# Plasticity-led evolution as an intrinsic property of developmental gene regulatory networks

**DOI:** 10.1101/2023.05.25.542372

**Authors:** Eden Tian Hwa Ng, Akira R. Kinjo

## Abstract

The Modern Evolutionary Synthesis seemingly fails to explain how a population can survive a large environmental change: the pre-existence of heritable variants adapted to the novel environment is too opportunistic, whereas the search for new adaptive mutations after the environmental change is so slow that the population may go extinct. Plasticity-led evolution, the initial environmental induction of a novel adaptive phenotype followed by genetic accommodation, has been proposed to solve this problem. However, the mechanism enabling plasticity-led evolution remains unclear. Here, we present computational models that exhibit behaviors compatible with plasticity-led evolution by extending the Wagner model of gene regulatory networks. The models show adaptive plastic response and the uncovering of cryptic mutations under large environmental changes, followed by genetic accommodation. Moreover, these behaviors are consistently observed over distinct novel environments. We further show that environmental cues, developmental processes, and hierarchical regulation cooperatively amplify the above behaviors and accelerate evolution. These observations suggest plasticity-led evolution is a universal property of complex developmental systems independent of particular mutations.

## Introduction

According to the Modern Evolutionary Synthesis, the standard theory of evolution, all possible phenotypic variation is almost purely explained by genetic variation^1^, either ignoring environmental contributions or treating them as noise^2, 3^. In this sense, the standard theory is said to be a theory of mutation-led evolution. Therefore, the only means for an individual to survive a large environmental change is to possess mutations that produce a phenotype already adapted to the novel environment. However, natural selection selects adaptive phenotypes in the current environment, making the pre-existence of phenotypes adapted to novel environments highly unlikely^4^. Suppose instead that adaptive variants only appear after the environmental change. In that case, adaptation requires searching for new adaptive mutations, which is likely too slow for the population to survive^5^.

Phenotypic plasticity, the ability to change the expressed phenotype in response to environmental cues, has been proposed to remedy the above problem because it could produce a phenotype with higher fitness in a novel environment without a change in the genotype. Phenotypic plasticity arises from the developmental process, which integrates genetic and environmental information to generate a phenotype^6–8^. Biological experiments suggest that formerly conditionally expressed traits can become constitutively expressed through a process called genetic assimilation^9^. Genetic assimilation was later generalized to genetic accommodation to include any adaptive refinement of phenotype regulation^6, 10^.

In *plasticity-led evolution*, the novel adaptive phenotype is initially induced by novel environmental cues. If the novel environment is persistent, then the novel phenotype undergoes genetic accommo- dation^6, 11, 12^. This has been deemed to resolve the problem of gradualism implied by mutation-led evolution^13^. Levis and Pfennig proposed the following four criteria for plasticity-led evolution^11^:

1. The novel adaptive phenotype is initially induced by plasticity.
2. Cryptic mutations are uncovered as a result of the plastic response.
3. The phenotype undergoes a change in regulation.
4. The phenotype undergoes adaptive refinement under selection.

There are numerous studies on natural populations to evaluate whether plasticity-led evolution has occurred^11^. However, the mechanisms by which plasticity-led evolution is made possible remain unclear, resulting in misunderstanding and confusion on the meaning and implications of plasticity-led evolu- tion^1, 10, 12, 14^. For example, if plastic responses are regulated by particular genes, plasticity itself is a heritable trait^1^ and, hence, is subject to natural selection. The “plasticity-led evolution” based on such gene-regulated plasticity is, therefore, still within the framework of Modern Evolutionary Synthesis, and hence the problem of gradualism remains. This suggests that plasticity in plasticity-led evolution should emerge from non-genetic causes^5, 13, 15^. This is not to say that plasticity does not depend on genetic information. It is to say that the plasticity in plasticity-led evolution should be an emergent, collective property of the developmental system, including the genome and environment as a whole^5, 13, 15, 16^. A similar phenomenon has been suggested for evolutionary capacitance^17^.

To pursue this possibility, we study computational models of evolutionary developmental systems. Such models should be able to express changes in phenotype in response to changes in environmental cues and possess a developmental framework and hierarchical regulation^18, 19^. In plasticity-led evolution, the environment encountered by a population plays two roles. One induces phenotypic response (environment- as-inducer), and the other selects phenotypes more adapted to it (environment-as-selector). Existing works incorporate some aspects of phenotypic response to different environmental cues^20–23^. However, so far, there seem to be almost no studies that explicitly correlate the two roles of the environment^4^. A notable exception is Draghi and Whitlock,^24^, who used correlated environment-as-selector and environment- as-inducer (modeled as 2-dimensional vectors) to study the effect of genetically encoded plasticity on adaptation. The developmental process is often overlooked in traditional quantitative genetics models as it does not directly contribute to phenotypic variation^3, 25, 26^. However, it is an essential process that integrates environmental and genetic information into phenotype^6, 7^. Developmental processes are naturally modeled by gene regulatory networks (GRNs)^27, 28^. Biological GRNs have a hierarchical organization^18, 19, 28^. However, most studies of computational models ignore this biological aspect of development (except for Xue et al.^29^, but their model lacks developmental regulation).

We demonstrate that GRN models incorporating all the above ingredients can satisfy the Levis- Pfennig criteria for plasticity-led evolution under large environmental changes^11^. We also illustrate how these ingredients cooperatively enhance adaptation and accelerate evolution. We further show that this model exhibits plasticity-led evolution as a generic feature independent of specific mutations or particular environmental changes.

## Modeling

A computational model of plasticity-led evolution should incorporate several core notions: environment (-as-inducer and -as-selector), gene regulatory network (GRN), developmental process, selection, and reproduction. The environment-as-inducer represents the role of the environmental cue in determining phenotype alongside the genome. The environment-as-selector represents the role of the environment as a selection agent. In our work, we assume that these two roles of the environment are highly correlated. The GRN represents the regulation of phenotype expression through gene-gene and gene-environment interactions. We model the developmental process as the recursive regulation of gene expressions over time to express the phenotype. Selection favors individuals that have adult phenotypes that better match the environment-as-selector. Selected individuals then reproduce. Their genomes are recombined and mutated to produce the next generation of individuals.

A minimal model incorporating all these notions is a recursive GRN introduced by A. Wagner^27^. In the Wagner model, the gene expression at the *s*-th stage of development is represented by a vector *g*(*s*). The genome is represented by a matrix *G* where the (*i, j*) element represents the regulatory effect of the *j*-th gene on the *i*-th gene. The recursive equation defining the developmental process of the Wagner model is given by

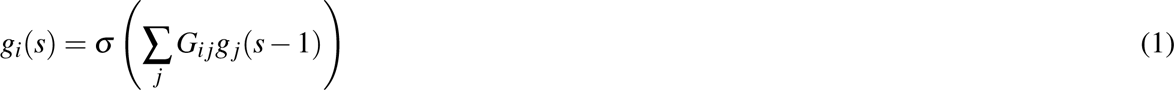

where *σ* is an activation function.

The developmental process is naturally represented by the sequence of vectors *g*(0)*, g*(1)*, g*(2)*,···* . The individual’s phenotype is usually taken as the steady state of Eq.(1) if it converges. The Wagner model has been used to demonstrate the evolution of mutational robustness^27^, evolutionary capacitance^17^, the link between mutational and environmental robustness^30^, the role of robustness in evolution^31, 32^, the role of phenotypic plasticity in directing evolution^33^ and the emergence of bistability^22, 23^. These works provide indirect evidence that the Wagner model has the potential to exhibit plasticity-led evolution^4^.

However, most previous works using the Wagner model did not include correlations between the environment-as-inducer and environment-as-selector to validate whether a plastic response is adaptive.

These works also do not emphasize the roles of developmental processes or hierarchical regulation of GRNs. We, therefore, extend the Wagner model by introducing these extra features.

### Macro-environment and environmental cues

Recall that the environment plays two roles in eco-evo-devo biology. The “environment-as-inducer” forms a cue integrated into phenotype expression. The “environment-as-selector” determines the fitness of adult individuals. We generalize the idea of Draghi and Whitlock^24^ to a higher-dimensional case to model a wider variety of environments and phenotypes. We define the macro-environment as a 200-dimensional vector **e** representing the average environment exerted on the population. Each element of **e** takes a+1 or *−*1 value. We modeled each individual’s environmental cue *e* as the macro-environment **e** with noise by randomly flipping 5% of elements of **e**. This environmental cue *e* may be considered as the micro-environment of an individual.

### Genome and developmental process

We now introduce several variants of the Wagner model. The *Full* model (Fig. 1**a**) is the main focus of this study, which incorporates response to environmental cues, developmental process, and hierarchical regulation. As controls, we also introduce *NoHier*, *NoCue*, and *NoDev* (Fig. 1**b**-**d**) models to highlight the role of each ingredient in plasticity-led evolution.

**Figure 1.**
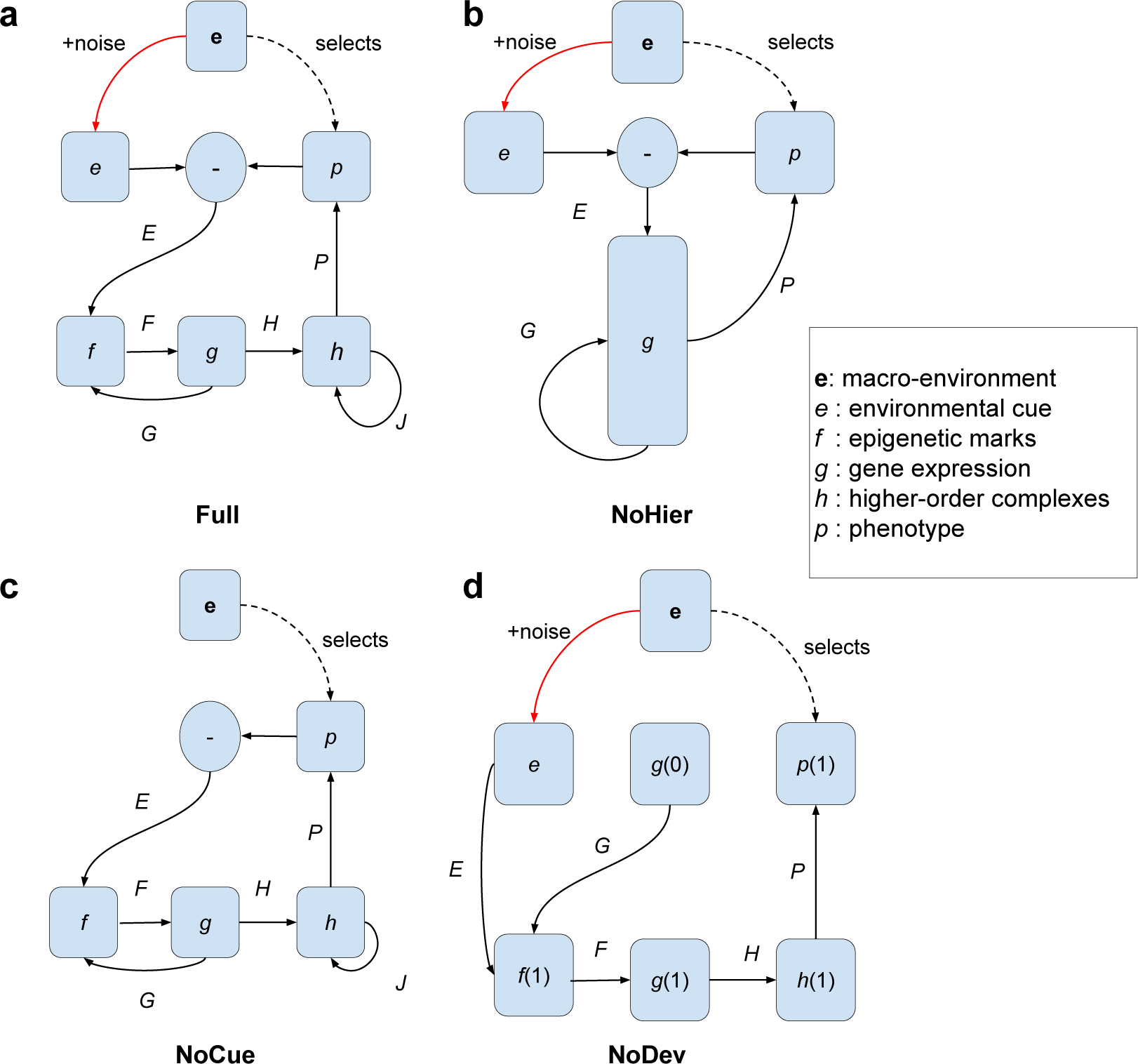
Diagram representing regulation between different layers of different models. Boldface **e** represents the macro-environment. *e, f, g, h, p* represent vectors. Black solid arrows represent regulatory interactions. The red solid arrow represents noise. The black dotted arrow represents selection. *E, F, G, H, J, P* represent regulatory matrices. **(a)** Full model (Eq.. 2); **(b)** NoHier (Eq. 3); **(c)** NoCue (Eq. 4); **(d)** NoDev (Eq. 5).

### Full Model

We introduce a vector *p*(*s*) representing the phenotype expressed at the *s*-th stage of development and *p̃*(*s*) as the exponential moving average of the phenotype (see **Methods**). To reflect the hierarchical regulation of GRN elements (such as epigenetic marks, RNAs, and proteins), we further introduce a layer of vector *f* (*s*) to represent epigenetic marks and a layer of vector *h*(*s*) to represent higher-order complexes (such as proteins, supramolecular complexes, etc.). Thus, we assume the following mutually recursive equations:

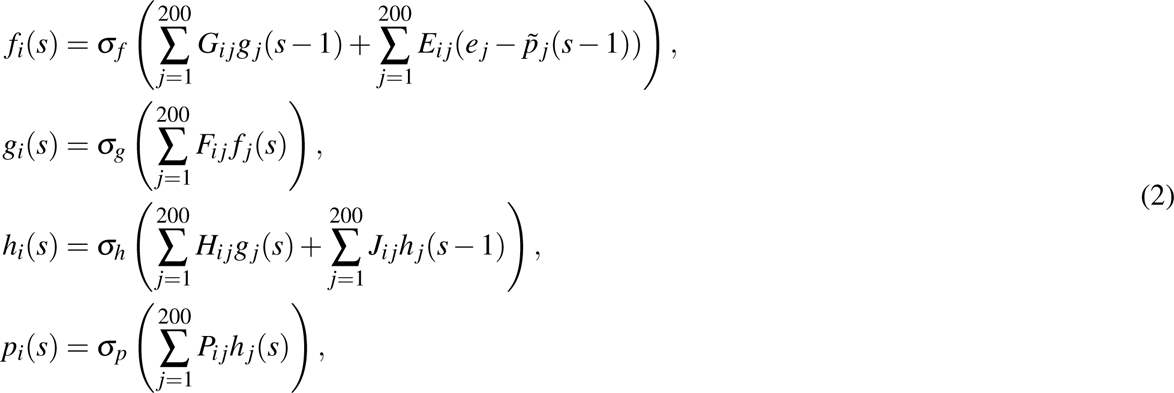

where *E* is a matrix that represents the environmental regulation of epigenetic marks, *F* is a matrix representing the epigenetic regulation of gene expression levels, *G* is a matrix representing the genetic regulation of epigenetic marks, *H* is a matrix representing the genetic regulation of higher-order complexes, *J* is a matrix representing interactions among higher-order complexes, *P* is a matrix representing regulation of the phenotype. Therefore, the matrix ensemble *{E, F, G, H, J, P}* represents the individual’s genome.

We set the base matrix density as *ρ*_0_ = 0.02, which is the density of each matrix in the Full model. The matrix densities for the other models are determined such that the number of nonzero elements is equal between different models on average. *σ_f_, σ_g_, σ_h_,* and *σ_p_* are activation functions based on arctangent or hyperbolic tangent functions (see **Methods**). The initial conditions are set to *f* (0) = **0**, *g*(0) = **1**, *h*(0) = **0**, and *p*(0) = **0**, where **0** is the zero vector and **1** is the vector with all elements equal to 1. *f* (*s*)*, g*(*s*)*, h*(*s*), and *p*(*s*) are all 200-dimensional vectors in the Full model, the values of their elements can be interpreted as their respective normalized values. We iteratively compute the state vectors *f* (*s*)*, g*(*s*)*, h*(*s*), and *p*(*s*) for *s* = 1, 2*, · · ·* until the phenotype *p*(*s*) converges (see **Methods**).

Note that the environmental cues are fed to the system as *e− p̃*(*s −* 1) rather than *e* alone. In reality, no environmental cues can directly influence an organism, but only through some receptors or sensors, which are part of the phenotype^34^. The *e− p̃*(*s −* 1) term can represent these cue-receptor/sensor interactions most straightforwardly. As the population adapts towards **e**, (selected elements of) *p* approaches **e**, and their difference converges to **0**. In other words, the influence of the environmental cues on the adult phenotype decreases as adaptedness increases. In this way, we expect to model genetic assimilation^13^. In contrast, early developmental stages (e.g., embryos) are more strongly influenced by environmental cues, in line with experimental observations^9, 35, 36^.

### NoHier model

To study the effect of hierarchical structure on GRNs, we introduce a developmental model without a hierarchical structure, which we name the *NoHier* model (Fig. 1**b**):

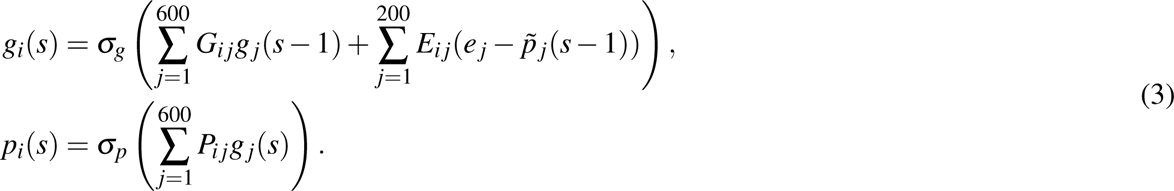

To preserve the degrees of freedom between the Full and NoHier models, the vector representing gene expression levels *g*(*s*) is set to 600-dimensional for the NoHier model. The matrix densities of the NoHier model are adjusted so that the number of non-zero elements is the same as the Full model, on average (Table 1). The dimensions of vectors representing the environment *e* and phenotype *p*(*s*) are kept at 200.

**Table 1.** Summary of parameters in model variants. dim *g* is the dimension of the *g* vector. All other vectors, if present, are 200-dimensional. Matrix densities (*ρ_G_, ρ_P_, ρ_E_, ρ_H_*) are chosen to ensure that the number of non-zero elements is equal between different models on average. Our study used *ρ*_0_ = 0.02. *ω_f_, ω_g_, ω_h_* and *ω_p_* are scaling constants for the input of the respective activation functions (squared values are shown; see Methods).

|  | Full | NoHier | NoCue | NoDev |
| --- | --- | --- | --- | --- |
| vectors | $e - p, f, g, h, p$ | $e - p, g, p$ | $-p, f, g, h, p$ | $e, f, g, h, p$ |
| matrices | $E, F, G, H, J, P$ | $E, G, P$ | $E, F, G, H, J, P$ | $E, F, G, H, P$ |
| $\dim g$ | 200 | 600 | 200 | 200 |
| $\rho_G$ | $\rho_0$ | $4\rho_0/9$ | $\rho_0$ | $\rho_0$ |
| $\rho_P$ | $\rho_0$ | $\rho_0/3$ | $\rho_0$ | $\rho_0$ |
| $\rho_E$ | $\rho_0$ | $\rho_0/3$ | $\rho_0$ | $\rho_0$ |
| $\rho_H$ | $\rho_0$ | N.A. | $\rho_0$ | $2\rho_0$ |
| $\omega_f^2$ | $3 \times 200\rho_G$ | N.A. | $2 \times 200\rho_G$ | $2 \times 200\rho_G$ |
| $\omega_g^2$ | $200\rho_G$ | $600\rho_G + 2 \times 200\rho_E$ | $200\rho_G$ | $200\rho_G$ |
| $\omega_h^2$ | $2 \times 200\rho_H$ | N.A. | $2 \times 200\rho_H$ | $200\rho_H$ |
| $\omega_p^2$ | $200\rho_P$ | $600\rho_P$ | $200\rho_P$ | $200\rho_P$ |

### NoCue model

To study the effect of environmental cues on development, we introduce a developmental model where the environmental cue *e* is absent, which we name the *NoCue* model (Fig. 1**c**):

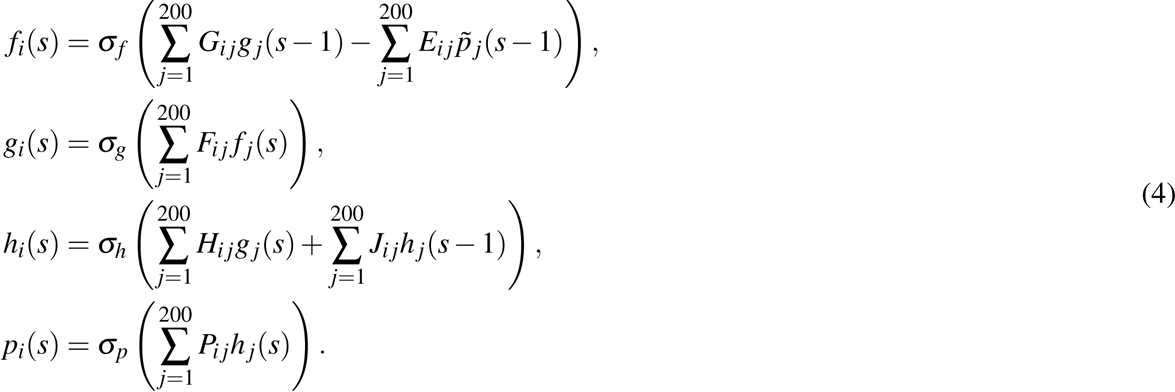

Apart from the absence of the vector *e*, this is identical to the Full model.

### NoDev model

To highlight the importance of the developmental process, we use the Full model described by Eq. (2) but set the maximum number of developmental steps to 1 for minimal development. We name this the *NoDev* model (Fig. 1**d**):

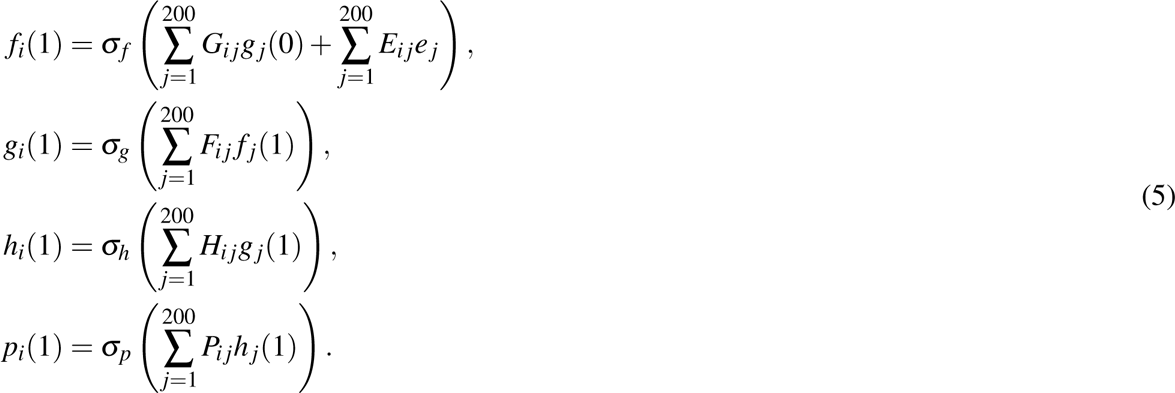

Hence, *p*(1) is immediately considered the adult phenotype. To compensate for the absence of self-regulation of the *h* layer, we double the density of the *H* matrix (Table 1).

### Natural selection and reproduction

We evaluate each individual’s fitness by matching the macro-environment **e** with the adult phenotype *p* (assuming that *p* converges). In other words, the macro-environment **e** plays the role of environment- as-selector by being the optimal phenotype. We also included the number of developmental steps up to convergence into the fitness calculation such that individuals with fewer developmental steps are favored. We define the raw fitness of the *i*-th individual as:

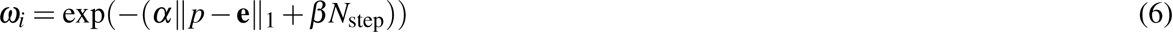

Where ||*p−* **e**||_1_ is the absolute (L1) distance between the first 40 (out of 200) elements of the adult phenotype *p* and the corresponding elements of the macro-environment **e**, *N*_step_ is the number of developmental steps until convergence, and we set *α* = 20 and 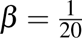. In other words, only 40 traits (elements of the phenotype vector) are subject to selection, and other 160(= 200 *−* 40) traits are allowed to evolve freely. In this paper, we call the value of ||*p−* **e**||_1_ mismatch. Individuals whose phenotype *p*(*s*) does not converge before a pre-specified number (200) of steps are given a fitness value of zero. The population in which all individuals have completed the developmental process is called the adult population in the following. The relative fitness of the *i*-th individual of the adult population is given by

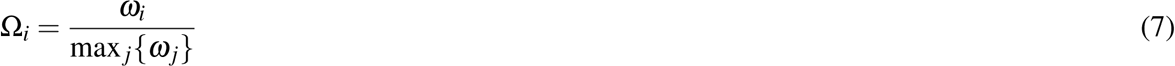

where max *_j_{ω_j_}* is the maximum raw fitness among the adult population.

The offsprings of individuals are generated as follows:

1. Initialize the selected population as an empty set.
2. Uniformly sample individual *i* from the adult population.
3. Sample a random number *r* from the uniform distribution between 0 and 1.
4. If *r <* Ω*_i_*, then a copy of the *i*-th individual is added to the selected population.
5. The *i*-th individual is put back to the adult population.
6. Repeat steps 2-5 until the size of the selected population reaches the maximum population size of 1000 individuals.
7. Individuals *i* and *i* + 1 where *i* = 1, 3, 5*, · · ·* of the selected population are paired to become parents.
8. For each pair of parents, two offsprings are produced by randomly shuffling the corresponding rows of the genome matrices with probability 0.5 between the parents.
9. Duplicate the offspring population. Now we have two populations of 1000 offsprings each.
10. The genome matrices of both offspring populations are independently mutated with a given probability (see **Methods**).

We then develop one offspring population in the “ancestral” environment and the other in the “novel” environment (see the following subsection for definitions of the ancestral and novel environments). Only the individuals in the “novel” environment are subject to selection. The population in the “ancestral” environment is used only for comparison and is discarded after measurement. Adaptive evolution of a population in an environment is therefore modeled by repeated cycles of development, selection, and reproduction under the “novel” environment.

### Simulating plasticity-led evolution

To study plasticity-led evolution, we require at least two environments: the ancestral environment, to which the population is adapted, and the novel environment, to which the population is to adapt. To let the population adapt to an environment, we simulate adaptive evolution for 200 generations. We call this duration an *epoch*, which is considered a unit of evolutionary time scale. In each epoch, the macro-environment is set constant. For each model, we simulated adaptive evolution for 50 epochs in turn. Between two consecutive epochs, we introduce a large environmental change by randomly flipping 50% of the elements of the macro-environment vector **e**. The first 40 epochs serve as the “training phase,” where the randomly initialized population is equilibrated. In addition, we expect the developmental systems to learn how to respond to large environmental changes during this phase. We then tracked the final ten epochs to assess the properties concerning the criteria of plasticity-led evolution. In each epoch, the last adapted and current environments are regarded as the ancestral and the novel environments, respectively.

## Results

We first present a visualization of the trajectory of the phenotype and genotype over evolution. By looking at the initial phase of the trajectory, we assess plastic response and the uncovering of cryptic mutations. The visualization also lets us track adaptive change in mean value and variation of genotype and phenotype over evolution. We compared different models in light of the Levis-Pfennig criteria of plasticity-led evolution^11^. We also present additional results closely related to plasticity-led evolution.

### Visualizing evolution: The Genotype-Phenotype plot

To visualize evolution, we plotted the phenotypic value in the ancestral and novel environments against the genotypic value of the population over evolution for each epoch. We call this plot the *genotype-phenotype plot* (Fig. 3). Here, the phenotypic value is computed by projecting the phenotype vector onto an axis where 0 corresponds to the adult phenotype perfectly adapted to the ancestral environment and 1 to that perfectly adapted to the novel environment. Similarly, the genotypic value is computed by projecting the genome matrices (construed as a vector) onto an axis where a value of 0 corresponds to the average genotype of the first generation and a value of 1 to the average genotype of the 200th generation (see **Methods**). Each generation’s projected phenotypes and genotypes are those after development but before selection. The trajectory of adaptive evolution in the novel environment generally proceeds from the lower left corner to the upper right corner in the genotype-phenotype plot.

**Figure 2.**
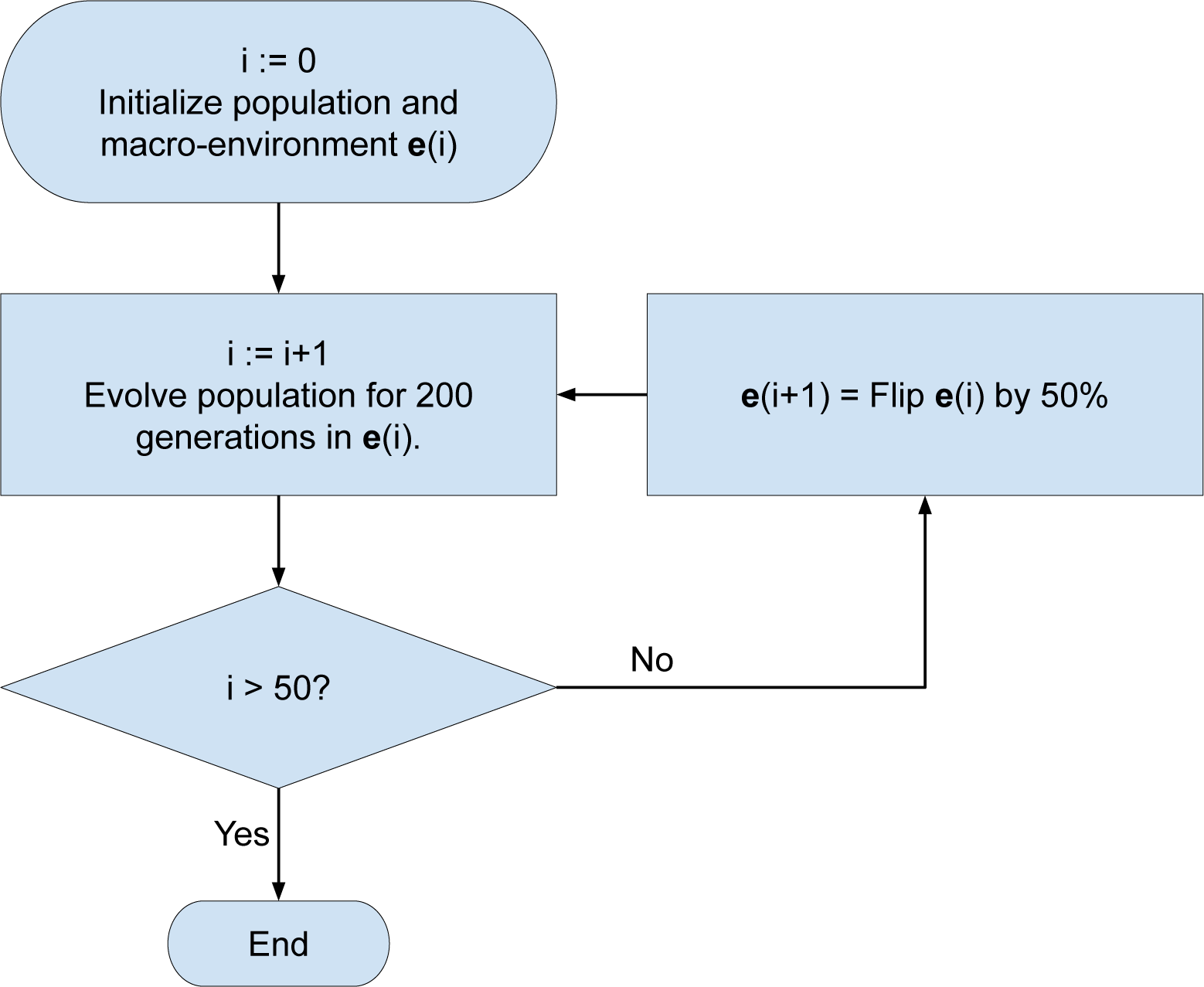
Simulating plasticity-led evolution. During each epoch, a population of individuals is subject to selection under a constant macro-environment. Each epoch lasts for 200 generations. At the end of each epoch, the macro-environment is changed by randomly flipping 50% of the environmental factors. This process is repeated so that the population is subject to selection under 50 different environments in turn. The first 40 epochs are the training phase; we tracked the final ten epochs to assess plasticity-led evolution.

**Figure 3.**
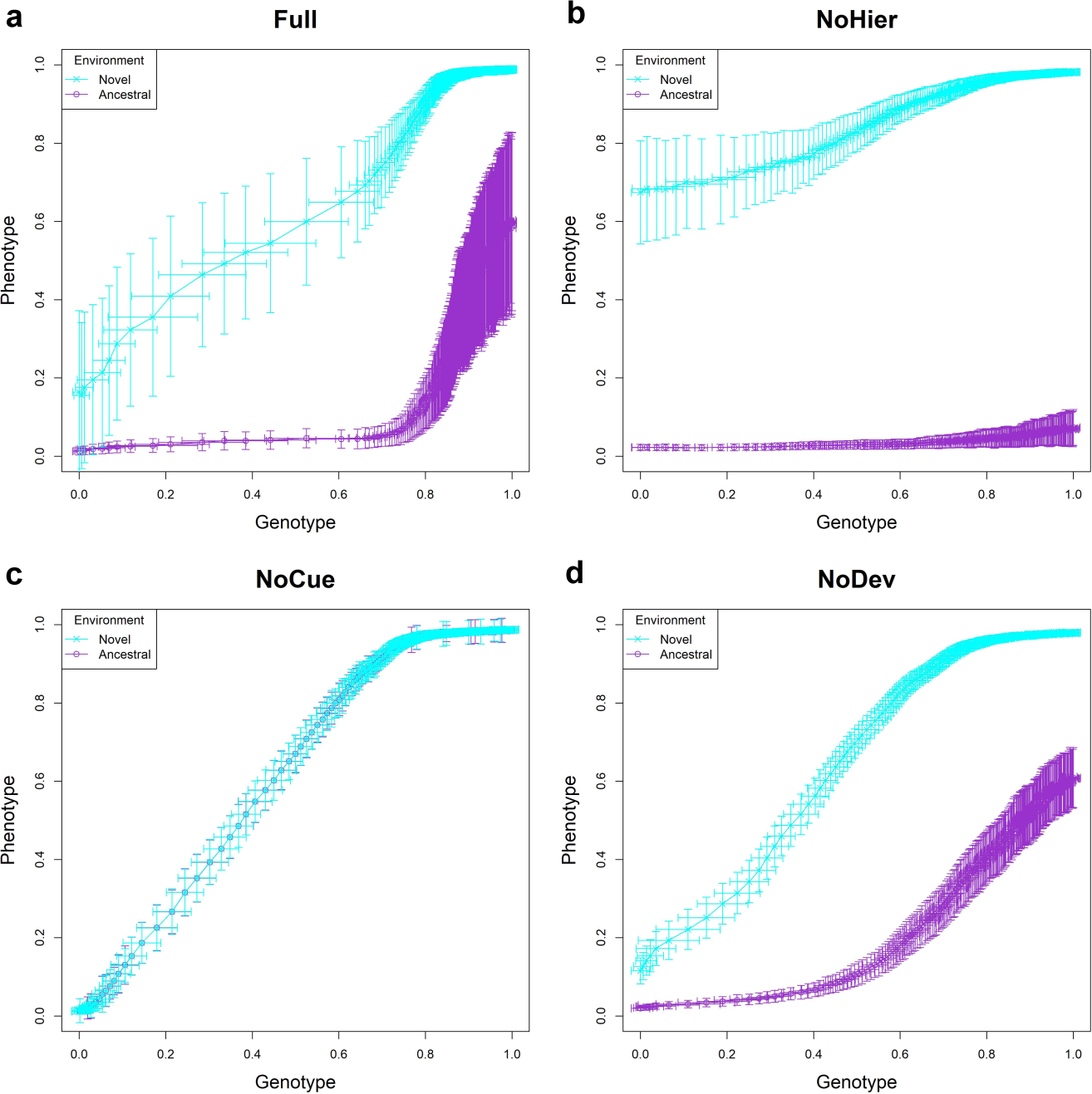
Trajectory of projected phenotype against projected genotype. The trajectory of one arbitrary epoch is shown. Each point represents the population average of genotypic value (horizontal axis) and phenotypic value (vertical axis) after development but before selection at each generation (error bars represent respective standard deviations). Cyan and purple represent populations in novel and ancestral environments, respectively. Projected phenotypic values of 0 and 1 correspond to perfectly fit phenotypes in ancestral and novel environments, respectively. Projected genotypic values of 0 and 1 correspond to the population average genome at first and 200th generations, respectively. Hence, the trajectory generally proceeds from the lower left to the upper right corner. **(a)** Full model; **(b)** NoHier; **(c)** NoCue; **(d)** NoDev. See also Supplementary Information.

From the genotype-phenotype plots, we observe that all discussed models are capable of adaptive evolution. The projected value of the novel phenotype tends to increase as the projected value of the genotype increases. The final average value of the projected phenotype for all models is about 1 in the novel environment after an epoch, indicating adaptation.

For the Full and NoHier models (Fig. 3 **a**,**b**), the phenotypes developed in the novel environment show substantially larger projected values than those developed in the ancestral environment in the first generation, indicating an adaptive plastic response. The standard deviation in phenotype in the novel environment is much larger than that in the ancestral environment in the first generation. This can be due to uncovered cryptic mutations or amplified phenotypic response to environmental noise. However, a higher rate of change of projected genotype observed in the Full and NoHier models during the early stages of adaptation indicates rapid purification of heritable variation during that phase. This suggests that the large standard deviation is more likely due to uncovered cryptic mutations (see subsection **Environmental and genetic variations induce correlated phenotypic variation** below). A notable difference between the Full and NoHier models is the much larger standard deviation in phenotype in the Full model after adaptation.

Environmental cues are absent in the NoCue model (Fig. 3 **c**). Consequently, there is no phenotypic plasticity, and the phenotypes expressed in the ancestral and novel environments are almost identical. The slight difference in phenotypes in the genotype-phenotype plot is purely due to the difference in mutations. The early stage of adaptation is very slow, as seen in the cluttered distribution of points in the lower left corner. This is followed by a rapid change in genotype, as seen in the sparse distribution of points around the center of the plot. This observation demonstrates that evolution in novel environments without phenotypic plasticity is characterized by an initial slow change followed by rapid adaptation once adaptive mutations appear.

We observe some adaptive plastic responses in the NoDev model (Fig. 3 **d**). However, on average, this adaptive plastic response is smaller than those of the Full or NoHier models. Similarly, there is less difference in the standard deviation in phenotypes between environments compared to the Full or NoHier models, indicating less uncovering of cryptic mutations. These observations highlight the importance of the developmental process in plasticity-led evolution.

We provide additional genotype-phenotype plots for different novel environments in Supplementary Information. We observe that these behaviors are consistent over different novel environments.

### Initial plastic responses tend to be adaptive

On the genotype-phenotype plots (Fig. 3), plastic responses are observed as a vertical shift between the ancestral and novel environments in the same generation. If the projected phenotype in the novel environment is greater than in the ancestral environment, then the plastic response is adaptive. We examined this shift in projected phenotype to detect the adaptive plastic response (Fig. 4 **a**). The Full and NoHier models exhibit large adaptive plastic responses on average. The NoDev model exhibits some adaptive plastic response, but it is less significant than those in the Full and NoHier models. As expected, the NoCue model does not show any plastic response.

**Figure 4.**
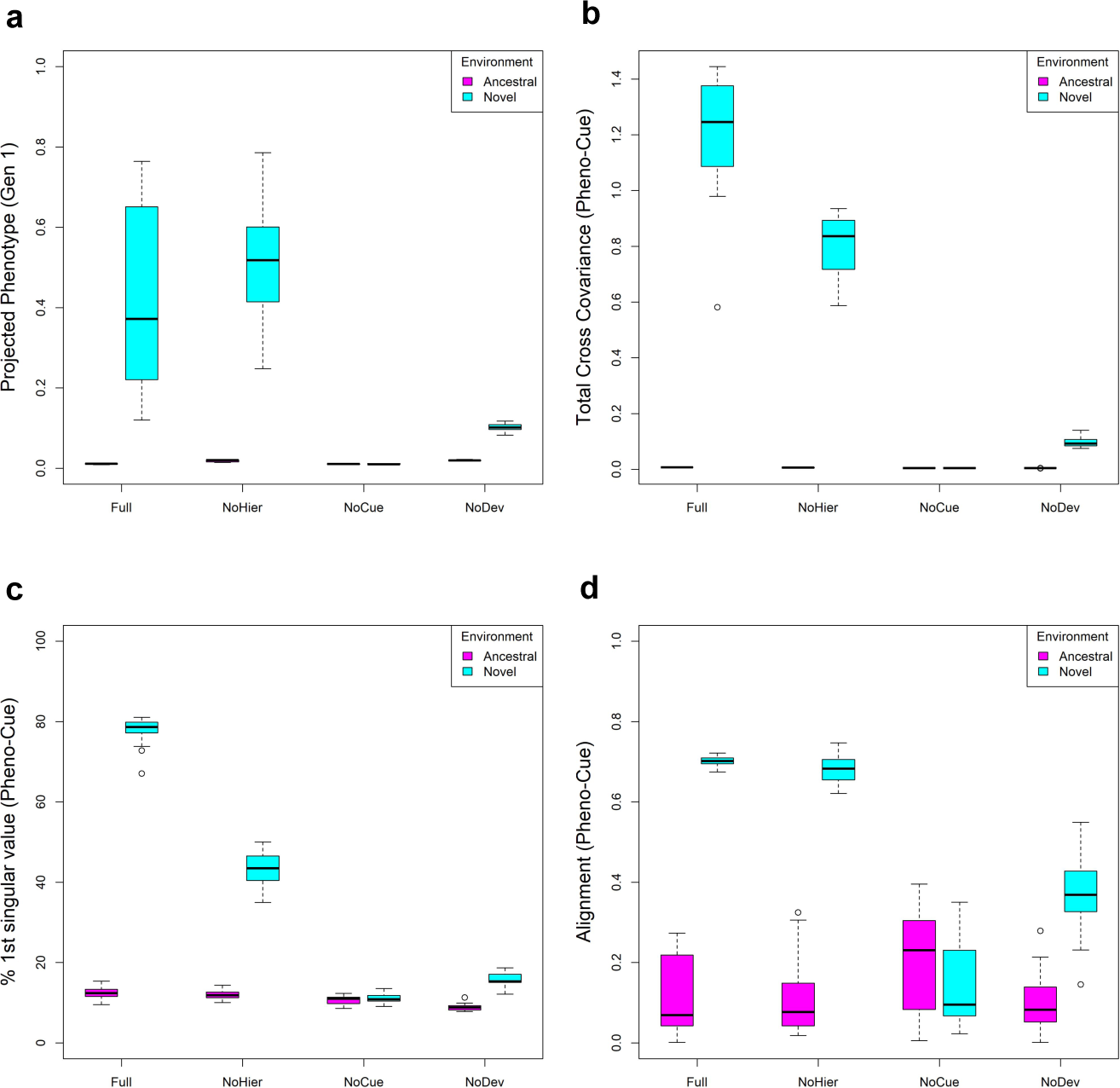
Adaptive plastic response in the first generation. (a) Boxplot of projected phenotype under ancestral and novel environment. **(b)** Boxplot of total cross-covariance between phenotype and environmental cue. **(c)** Boxplot of the percentage contribution of the first singular value of the cross-covariance matrix to the total cross-covariance between phenotype and environmental cue. **(d)** Boxplot of alignment between phenotype variation and environmental change, where the alignment is the correlation between the first phenotype singular vector of the Pheno-Cue cross-covariance matrix and the direction of environmental change.

Generally, the plastic response in the first generation is adaptive (except for the NoCue model), which shows that our models can learn to respond adaptively to a new environment from the past environments experienced in the training phase. This observation may appear surprising given that (1) the present macro- environment is uncorrelated with past environments and (2) environmental noise is uniformly distributed over environmental factors (i.e., no correlations between the factors, unlike “associative memory”^20, 21^). Nevertheless, the training process allows the models to learn each environmental factor independently of the other factors. As a result, the models can respond to each component of the environmental cues independently, albeit imperfectly.

To analyze the correlation between phenotype and environmental cues immediately after development but before selection in the novel environment, we computed their cross-covariance matrix in the first generation, which we call the Pheno-Cue cross-covariance matrix. We performed singular value decomposition (SVD) on the Pheno-Cue cross-covariance matrix to find the principal components (see **Methods**). The left singular vectors correspond to the principal axes of phenotypes (“phenotype singular vectors”), the right singular vectors correspond to the principal axes of environmental cues, and the singular values correspond to the cross-covariance between the corresponding left and right principal components. We performed this analysis for each model under ancestral and novel environments.

We observed a larger total cross-covariance between phenotypes and environmental cues in novel environments than in ancestral environments (Fig. 4 **b**). This suggests that populations are far more susceptible to environmental noise in novel environments than in ancestral environments. Among all models, the Full model has the largest cross-covariance, followed by the NoHier model. This suggests that a hierarchical structure in GRNs exaggerates phenotypic variation due to environmental noise. NoCue has little cross-covariance because the model itself is insensitive to environmental cues. NoDev exhibits smaller cross-covariances than the Full and NoHier models. This demonstrates that the developmental process amplifies variation in phenotype due to environmental noise in novel environments.

Previous works suggest that if the phenotypic variation is developmentally biased in line with the environmental change, such developmental bias, i.e., the tendency to generate certain phenotypes more readily than others, can facilitate evolution in novel environments^24, 37^. To detect any developmental bias in our models, we examined the proportion of the first singular component for each model under ancestral and novel environments. Fig. 4 (**c**) shows that the proportion of the first singular value tends to be larger in the novel environment than in the ancestral environment. This indicates that the phenotypic variation due to environmental noise is more biased in the direction of the first principal component in the novel environment. In particular, the Full model exhibits the greatest bias in phenotypic variation, where nearly 80% of the total cross-covariance is explained solely by the first singular component. The NoHier model shows less bias than the Full model, where the first singular component accounts for around 40% of the total cross-covariance. This shows that hierarchical regulation enhances developmental bias. The NoCue and NoDev models exhibit little bias in the novel environment. For the NoCue model, this is easily explained by the absence of environmental cues. For the NoDev model, the result emphasizes that the developmental process is essential in generating developmental bias. In contrast, all models exhibit small developmental biases in the ancestral environment. Together with the small total cross-covariances (Fig. b), this implies that all models are highly robust against environmental noise in adapted environments^31^. To see if the above developmental bias is aligned with environmental change, we computed the correlation between the first phenotype singular vector and the direction of environmental change (Fig. 4 **d**). The Full and NoHier models exhibit good alignment (roughly 0.7), with NoHier exhibiting a larger variance in alignment. Given the small proportions of the first singular values, the correlations for the NoCue and NoDev models are spurious.

### Developmental process uncovers cryptic mutations

As we mentioned above (**Visualizing evolution: The Genotype-Phenotype plot**), the large phenotypic variation in the novel environment compared to the ancestral environment during the early phase of evolution is most likely due to the uncovering of cryptic mutations (Fig. 3). We compared the magnitude of variance in projected phenotypes between ancestral and novel environments (Fig. 5 **a**). We observed a small variance in projected phenotype in every model in the ancestral environment. This can be explained by the evolved robustness in adapted environments^17, 22, 23, 27, 30, 31, 38^. The Full model exhibits the largest variance of projected phenotype in the novel environment on average, followed by the NoHier model, indicating that hierarchical regulation amplifies the effects of genetic variation. Compared to these, the NoDev model exhibits a smaller variance in projected phenotype in the novel environment. This indicates that the developmental process is critical in uncovering cryptic mutations. The NoCue model does not exhibit any difference in projected phenotype because environmental cues are necessary to uncover cryptic mutations.

**Figure 5.**
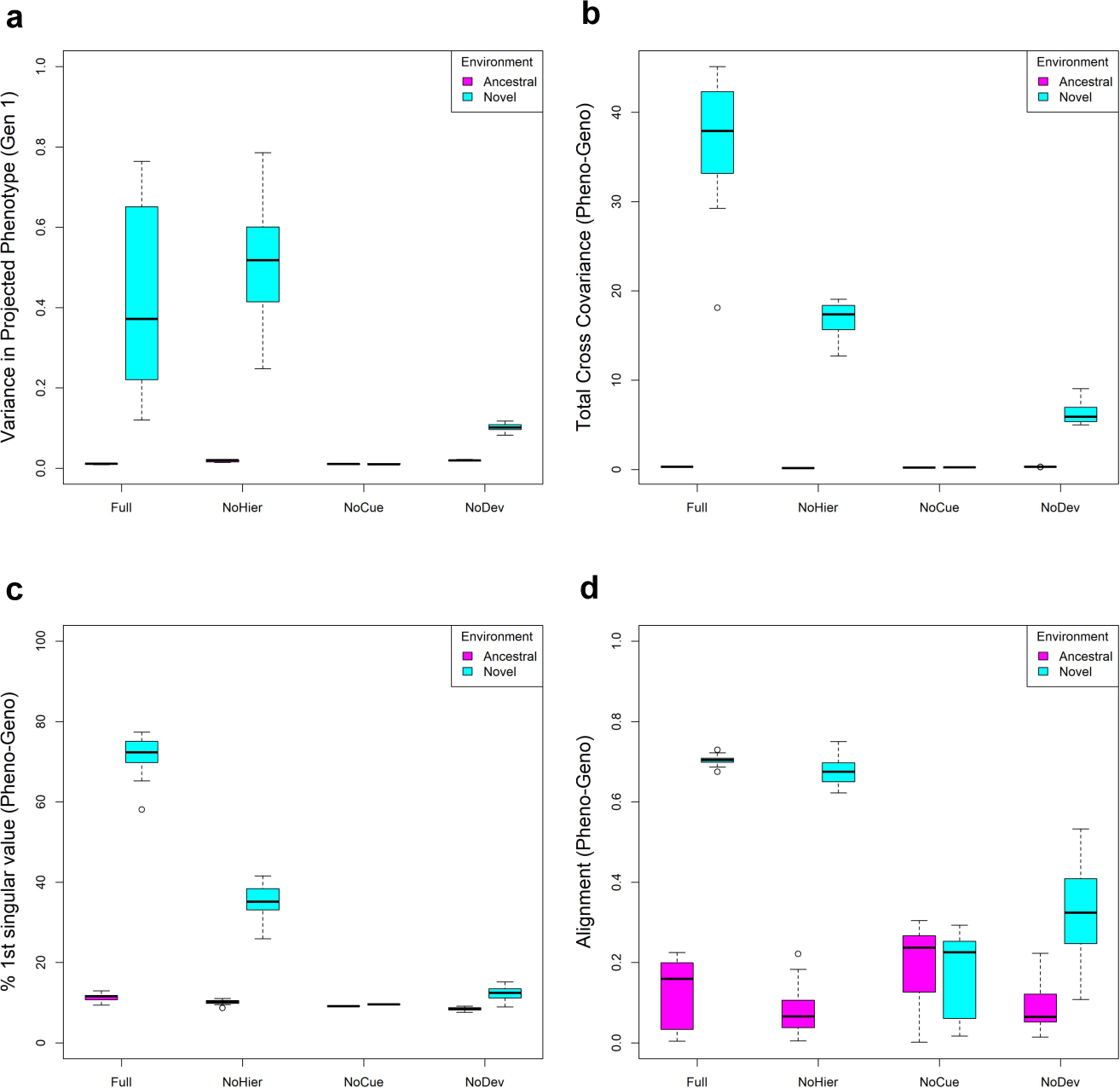
Uncovering of cryptic mutations. (a) Boxplot of variance in projected phenotype in ancestral and novel environments. **(b)** Boxplot of total cross-covariance (square of Frobenius norm) between phenotype and genome. **(c)** Boxplot of the percentage contribution of the first singular value of the cross-covariance matrix to total cross-covariance between phenotype and genome. **(d)** Boxplot of alignment between phenotype variation and environmental change, where the alignment is the correlation between the first phenotype singular vector of the Pheno-Geno cross-covariance matrix and the direction of environmental change. Variation due to mutations and environmental noise is qualitatively similar (c.f. Fig. 4).

To analyze the phenotypic variation due to mutations, we calculated the cross-covariance between the phenotype and the genome, called the Pheno-Geno cross-covariance matrix (see **Methods**). As we did for the Pheno-Cue cross-covariance matrix in the previous subsection, we performed SVD analysis on the Pheno-Geno cross-covariance matrix. Here, the left singular vectors still correspond to the principal axes of phenotypes (“phenotype singular vectors”), but the right singular vectors correspond to the principal axes of mutations instead of environmental cues.

We notice that the SVD analysis on the Pheno-Geno cross-covariance (Fig. 5**b**-**d**) is qualitatively similar to that of the Pheno-Cue cross-covariance (Fig. 4**b**-**d**). That is, the Full and NoHier models exhibit large total cross-covariances between phenotype and mutation (Fig. 5**b**), their phenotypic variation due to mutations are highly biased in novel environments (Fig. 5**c**), and phenotypic variations are biased in the direction of environmental change (Fig. 5**d**). On the other hand, NoDev and NoCue models have smaller total cross-covariance (Fig. 5**b**) and lack developmental bias (Fig. 5**c,d**).

### Environmental and genetic variations induce correlated phenotypic variation

We observed that the results from SVD analysis of the Pheno-Cue cross-covariance matrix are qualitatively similar to that of the Pheno-Geno cross-covariance matrix, leading to similar conclusions (c.f. Figs. 4 and 5). To examine the correlation between the phenotypic variation due to environmental noise and that due to mutations, we compared the first singular values of the Pheno-Cue and Pheno-Geno cross-covariance matrices (Fig. 6**a**). The distribution of the first singular values over different models is qualitatively similar between the Pheno-Cue and Pheno-Geno cross-covariances. That is, the first singular values decrease in the order of Full, NoHier, NoDev, and NoCue models in the novel environments. In contrast, all models exhibit minimal first singular values in the ancestral environment. Quantitatively, the first singular values of the Pheno-Geno cross-covariance matrices (right vertical axis of Fig. 6**a**) are much greater than those of the Pheno-Cue cross-covariance matrices (left vertical axis of Fig. 6**a**). This is because the number of elements of the genome vector is much larger than that of the environmental cue vector.

**Figure 6.**
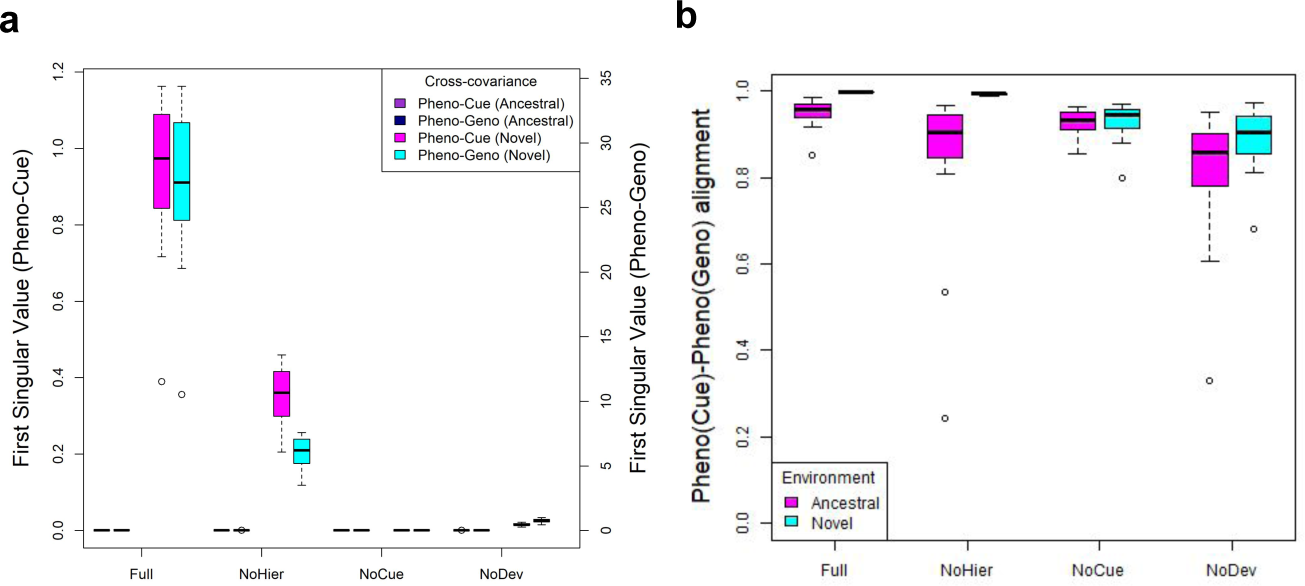
Environmental variation and genetic variation induce correlated phenotypic variation**. (a)** Boxplot of first singular values of Pheno-Cue and Pheno-Geno cross-covariance matrices in ancestral and novel environments. **(b)** Boxplot of (Pheno-Cue)-(Pheno-Geno) alignment; calculated as the correlation between the first phenotype singular vector of the Pheno-Cue cross-covariance matrix and the first phenotype singular vector of the Pheno-Geno cross-covariance matrix. These quantities were computed in the first generation after development but before selection in the novel environment.

We next calculated the correlation between the first singular vector of the Pheno-Cue cross-covariance matrix and that of the Pheno-Geno cross-covariance matrix to see the alignment between them (Fig. 6**b**). These axes are highly correlated, especially in the novel environment. In particular, the Full and NoHier models have almost perfect alignment in the novel environment. We remark that the apparent high correlations in all other cases are spurious because the phenotypic distributions of these populations are unbiased (c.f. low percentage of first singular values from Fig. 4**c** and 5**c**), and the spurious correlation is due to the idiosyncrasies of the SVD algorithm used rather than actual good alignment. These observations indicate the interchangeability of environmental and mutational perturbations in producing phenotypes, consistent with the existing literature^6, 30, 39, 40^.

### Full model exhibits fastest change in regulation

To study the change in regulation or the reorganization of the genome, we track the genetic variance (i.e., the sum of the variance in each element of the genome matrices) over evolution (Fig. 7**a**). A rapid decrease in genetic variance implies strong purifying selection. A gradual increase in genetic variance means the accumulation of neutral or beneficial mutations. The Full and NoHier models immediately experience strong selection in the novel environment, with the Full model experiencing the most stringent selection (largest decrease in the shortest time). The NoDev model experiences weaker selection than the Full and NoHier models. In contrast to the other models, the genetic variance of the NoCue model initially increases slightly and then decreases rapidly before gradually increasing again. The initial increase in genetic variation in the NoCue model may be attributed to the random search for adaptive mutations in the novel environment.

**Figure 7.**
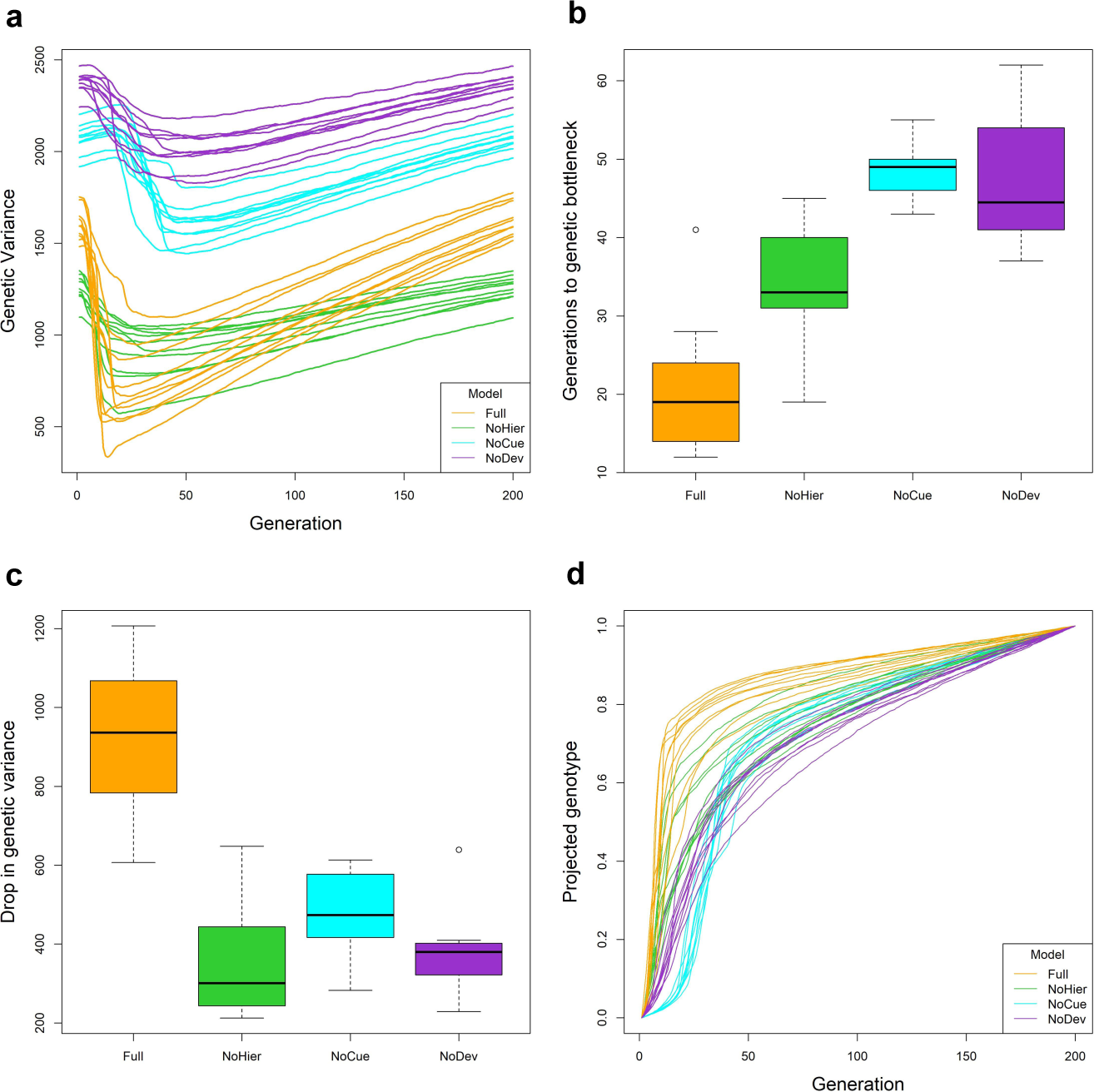
Change in regulation. (a) Trajectory of genetic variance over evolution. **(b)** Boxplot of the number of generations to genetic bottleneck, where the genetic bottleneck is the generation when the genetic variance is minimal (c.f. panel **a**). **(c)** Boxplot of drop in genetic variance between generation 1 and the genetic bottleneck. **(d)** Trajectory of projected genotype over evolution.

The genetic variance of the NoDev model is consistently greater than those of all the other models. Due to the lack of development, the NoDev model cannot fully express genetic variation in phenotype. Hence, selection cannot effectively purify the genetic variation. The genetic variance of the NoCue model is slightly smaller than that of the NoDev model but consistently greater than those of the Full and NoHier models. This suggests that most mutations remain latent in the NoCue model, highlighting the role of environmental cues in uncovering cryptic mutations.

We define the genetic bottleneck as the generation when the genetic variance is minimal (c.f. Fig. 7**a**). To track the rate of adaptation, we examined the number of generations to the genetic bottleneck measured from the first generation (Fig. 7**b**). The Full model has the lowest time, followed by the NoHier model. The NoDev and NoCue models reach the genetic bottleneck later than the Full and NoHier models.

We next examined the effectiveness of purifying selection by comparing the decrease in genetic variance from the first generation to the genetic bottleneck (Fig. 7**c**). The Full model exhibits the largest decrease in genetic variance among all models. The NoCue model exhibits a slightly larger decrease in genetic variance than the NoHier and NoDev models. This suggests that hierarchical developmental regulation enhances selection.

We also present the trajectory of projected genotypic value (c.f. Fig. 3) over evolution in the novel environment (Fig. 7**d**). The genotypic value generally increases rapidly when the genetic variance decreases rapidly, indicating strong purifying selection. On the other hand, the genotypic value generally increases slowly when the genetic variance increases slowly, indicating a gradual accumulation of neutral or adaptive mutations. These correlations indicate that the trajectory of the genetic variance is a suitable proxy for studying the change in regulation. Consistent with Fig. 7(**a**), we observe that the Full model exhibits the fastest increase (largest gradient) in genotypic value during the initial phase of evolution, indicating the fastest adaptation. For the NoCue model, the slow increase in genotypic value during the initial phase corresponds to the search for adaptive mutations via genetic drift.

### Full model exhibits greatest adaptive refinement

We now compare the effectiveness of adaptive refinement, or the increase in fitness in novel environments, between different models. To do so, we examined the trajectories of the mismatch between the adult phenotype and the selective environment for different models in the novel environments (Fig. 8**a**). All models exhibit a decrease in mismatch over evolution, trivially demonstrating that all models undergo adaptive refinement. All models, except for the NoCue model, exhibit a rapid decrease in mismatch immediately during the initial phase of evolution. In contrast, the mismatch of the NoCue model remains constant for around 25 generations before rapidly decreasing. This delay corresponds to the time required to find adaptive mutations in the novel environment (c.f. Fig. 7). This observation highlights the roles of environmental cues in inducing adaptive plastic phenotype and uncovering cryptic mutations to the selection, thereby accelerating evolution.

**Figure 8.**
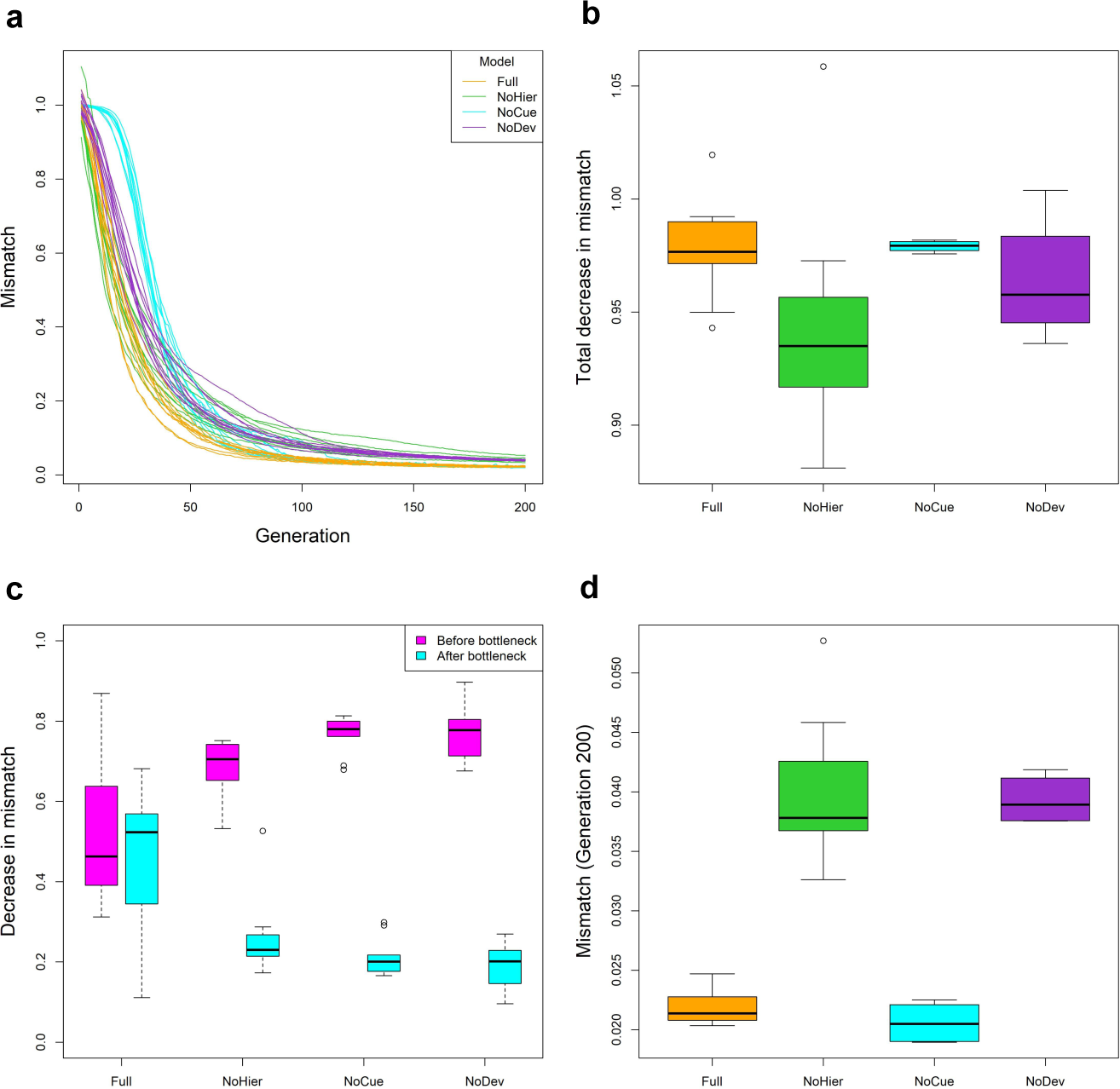
Adaptive refinement. (a) Trajectory of mismatch between phenotype expressed in a novel environment and selective environment over evolution. **(b)** Boxplot of the total decrease in a mismatch from Generation 1 up to Generation 200. **(c)** Boxplots of decrease in the mismatch before and after the genetic bottleneck. **(d)** Boxplot of the mismatch at generation 200.

When we compared the total decrease in mismatch among all models from Generation 1 to Generation 200, the Full and NoCue models exhibited greater decreases in mismatch than the NoHier and NoDev models on average (Fig. 8**b**). The decrease in mismatch for the NoCue model is the most consistent, while that for the NoHier model is the least consistent. For the NoHier model, the small decrease in mismatch could be attributed to low initial mismatch from the large adaptive plastic response (c.f. Fig. 4**a**).

We dissected the decrease in mismatch into the contributions before and after the genetic bottleneck (Fig. 8**c**; see also Fig. 7**a**). The Full model exhibits similar amounts of decrease in mismatch before and after the genetic bottleneck. In contrast, the decrease in mismatch before the genetic bottleneck tends to be significantly greater than after the genetic bottleneck for all other models. The implications of this behavior are unclear and could be an interesting topic for future studies.

To compare the quality of phenotypes after evolution, we examined the mismatch of the models at Generation 200 (Fig. 8**d**). The Full and NoCue models attain a significantly lower mismatch value at Generation 200 than the NoHier and NoDev models. This suggests that hierarchical developmental

### Full model accumulates the most mutations after the genetic bottleneck

The uncovering of cryptic genetic variation through plastic response under large environmental changes is one of the core criteria of plasticity-led evolution^11^. This criterion assumes that cryptic genetic variation has accumulated in the ancestral environment. We compare the accumulation of cryptic mutations between different models by measuring the increase in genetic variance after the genetic bottleneck in the novel environments (Fig. 9). The Full model exhibits a significantly larger increase in genetic variance than all the other models. In contrast, the NoHier and NoDev models exhibit the smallest increase in genetic variance on average.

**Figure 9.**
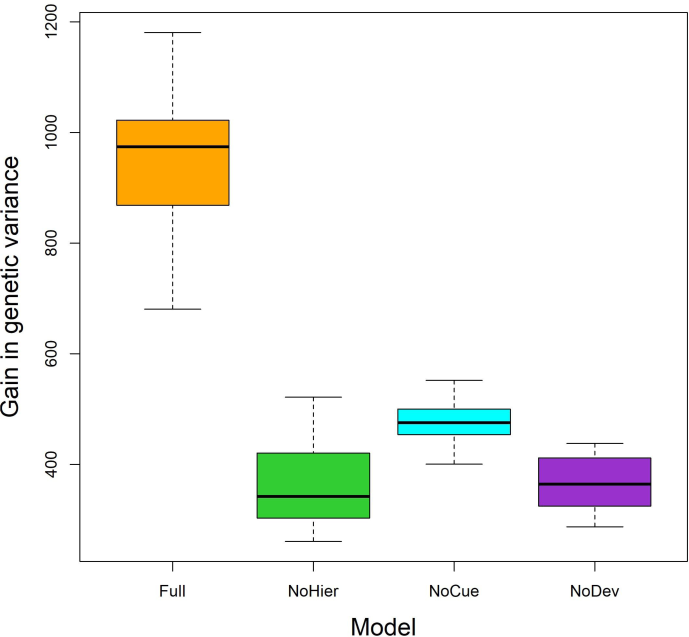
**Accumulation of cryptic mutations**. Boxplot of increase in genetic variance from genetic bottleneck up to generation 200. regulation is essential in refining phenotype quality. However, the lower mismatch for the NoCue model may also be explained by the absence of environmental noise in developmental regulation.

We also observe that the gain in genetic variance after the genetic bottleneck correlates to the drop in genetic variance before the genetic bottleneck (see Figs. 7 **a,b**). This can be explained by the fact that the length of each epoch is kept constant at 200 generations, so the same amount of cryptic mutations are accumulated in every epoch. Since the accumulation of cryptic mutations is made possible by the robustness of developmental systems^17, 31, 38, 41^, we may deduce that the Full model is the most robust of all the discussed models.

## Discussion

We have shown that the Full and NoHier models can satisfy all the Levis-Pfennig criteria of plasticity-led evolution under large environmental change. In particular, the Full model has additional favorable properties, such as amplifying the uncovering and the accumulation of cryptic mutations, accelerating change in regulation, and undergoing better adaptive refinement compared to the NoHier model. These observations suggest that environmental cues and the developmental process are essential for plasticity-led evolution, and hierarchical regulation enhances the desirable properties of plasticity-led evolution. These models consistently exhibit plasticity-led evolution over different environments, suggesting that plasticity-led evolution is an intrinsic behavior of these systems. This is not in line with the view that plasticity is explained by genetic variation^1^. We discuss the implications of this conflict below.

Contrary to our results, studies with natural populations suggest that plastic response is not always adaptive. For instance, spadefoot toad populations subjected to a dry condition during their larval stage express stunted development, potentially reducing fitness^42^. Another example is observed in populations of blue tits that use temperature cues to determine egg-laying periods: due to climate change, adult blue tits now prematurely lay their eggs, therefore, missing out on the optimal period when the caterpillar population (food source) is abundant^43^. In these studies, however, the “environment-as-inducer” does not match the “environment-as-selector,” so it is natural that the induced phenotypes are not adaptive. Ghalambor et al.^44^ introduced a population of guppies previously adapted to a high-predation (HP) environment to a low-predation (Intro) environment and compared the transcript abundance of the introduction populations with that of a population already adapted to a low-predation environment (LP). They claimed that transcription factors associated with initial plasticity are opposite to the direction of adaptive evolution, suggesting that non-adaptive plasticity can enhance evolution. However, the “non-adaptive plasticity” by Ghalambor et al.^44^ simply means that the genes responsible for the initial plastic response do not coincide with those responsible for later change of regulation, not that the plastic response is non-adaptive concerning the environmental change. In fact, Fig. 1 of Ghalambor et al.^44^ suggests that the plastic response of the Intro populations is indeed in the same direction as the LP population, hence adaptive.

Developmental bias, the tendency to generate certain phenotypes more readily than others, has been suggested as a critical mechanism for directing and thereby facilitating evolution^24, 37, 45^. We observed that phenotypic variation due to environmental cues is unbiased in adapted environments but is highly biased in novel environments for the Full and NoHier models (Fig. 4**c**). This biased phenotypic variation was consistently aligned with the direction of environmental changes (Fig. 4**d**). Furthermore, we observe the same behavior for phenotypic variation due to mutations (Figs. 5 **c**,**d**, and 6). These results imply that phenotypic variation due to uncovered mutations is aligned with environmental change, which, in turn, enhances the selection of adaptive mutations (and elimination of maladaptive mutations) in the novel environment. The exact causes of this behavior are unclear, it arises from the interplay between environmental cues and the developmental process since the NoCue and NoDev models do not exhibit this behavior.

Most studies on plasticity-led evolution observe the change in regulation through the changes in reaction norms^11, 16, 25, 46, 47^. However, reaction norms assume that phenotypic plasticity in the studied traits arises from particular genes that produce particular responses to particular environmental cues. Although reaction norms are helpful for studying the evolution of phenotypic plasticity^48^, this kind of phenotypic plasticity does not resolve the problem of gradualism implied by the Modern Evolutionary Synthesis. The problem may be resolved if, as proposed above, plasticity-led evolution is an emergent, collective property of the developmental system as a whole, independent of particular genetic variation. To validate this hypothesis, it may be helpful to study populations over many different novel environments and measure all traits subject to natural selection instead of some specific traits^12^.

## Methods

### Activation Functions

We use modified arctangent or hyperbolic tangent activation functions where the input, output, or both are scaled. For *σ_f_, σ_g_* and *σ_h_*, we use the following modified arctangent function:

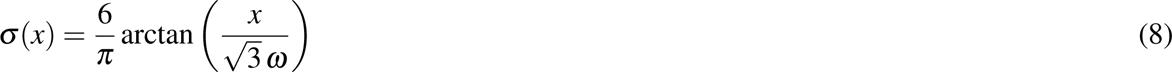

where the factors 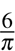 and 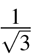 were derived in the same spirit as LeCun’s tanh function^49^. This maximizes the rate of change of *σ* (*x*) at around 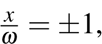, hence facilitating selection in the later stages of evolution.

The constant *ω* is introduced so that the estimated variance of 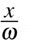 is 1 (See Table 1.) For *σ_p_*, we use the following hyperbolic tangent function:

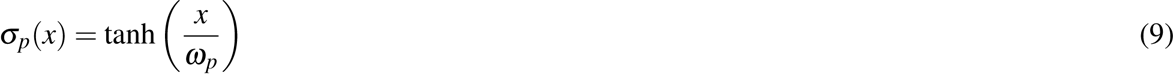

where *ω_p_* is a constant introduced so that the estimated variance of 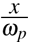 is 1 (See Table 1).

### Convergence of developmental process

To check the convergence of the developmental process of an individual, we used the limit of the exponential moving average (EMA) of the phenotype vector *p*(*s*). Denote the EMA of the phenotype as the vector *p̃*(*s*) and its variance as the vector *v*(*s*). Let 0 *< α <* 1 be given as the “step size” of the exponential moving average. In our work, we used *α* = 1*/*3. The values of *p̃*(*s*) and *v*(*s*) are recursively updated as follows:

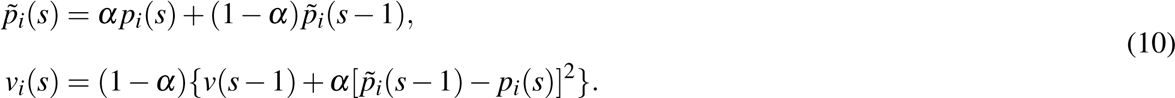

We say that the phenotype has converged when ∑*_i_ v_i_*(*s*) *<* 10*^−^*^5^ for *s ≤* 200 and take the adult phenotype as *p̃*_*i*_(*s*), otherwise, we say that the phenotype does not converge.

### Mutation

To randomly introduce mutations during reproduction, we let the mutation rate and matrix density be *γ* and *ρ*, respectively, between 0 and 1. The mutation rate *γ* represents the proportion of the genome matrix elements to be mutated per reproduction. We mutate the genome of each offspring as follows:

1. Initialize *n* = 0.
2. Sample the number of mutations *N* from a Poisson distribution where the mean is the product between *γ* and the genome size. (In our work, we used *γ* = 0.005, so we sampled from a Poisson distribution with mean *λ* = 0.005 *×* 200 *×* 200 *×* 6 = 1200.)
3. 3. Uniformly select an element in the genome matrix ensemble. The selected element is set to 0 with probability 1 *−ρ*, +1 with probability *ρ/*2, and *−*1 with probability *ρ/*2. Increase *n* by 1.
4. 4. If *n < N*, return to step 3. Otherwise, terminate the process.

### Visualizing evolutionary trajectory on genotype-phenotype space

To visualize the evolutionary trajectory of a population, we project the phenotype and genotype of individuals in the population at each generation onto a 2-dimensional genotype-phenotype space. First, the phenotype axis is defined as 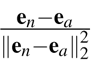where **e***_n_* is the first 40 elements of the novel macro-environment, **e***_a_* is the first 40 elements of the ancestral macro-environment and ||**e***_n_* − **e***_a_*||_2_ is the Euclidean (L2) distance between **e***_n_* and **e***_a_*. We consider only the first 40 out of 200 elements because they correspond to the traits subject to selection. We project the phenotype *p* of an individual as:

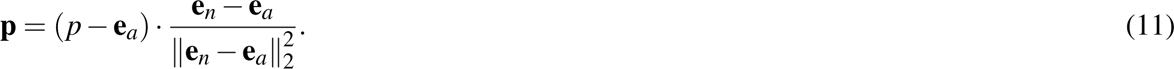

This way, the projected phenotypic values **p** of 0 and 1 correspond to phenotypes perfectly adapted to the ancestral and novel environments, respectively.

Next, the genotype axis is defined as follows. Let **G***_i j_* be the vectorized genotype matrix of the *i*-th individual of the *j*-th generation. Let 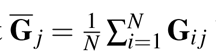 be the population average genotype vector on the *j*-th generation. The genotype axis is defined as 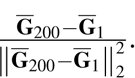 . We project the genotype of the *i*-th individual of the *j*-th generation onto a genotype axis as:

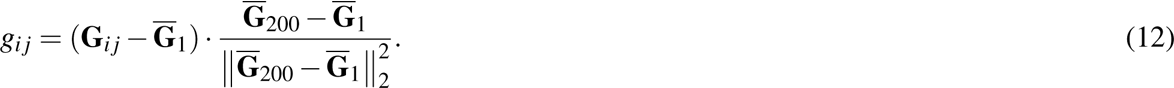

This way, projected genotypic values *g_i j_* of 0 and 1 correspond to the average genotypes before and after one epoch of evolution in a novel environment, respectively.

### Singular value decomposition (SVD) analysis of cross-covariance matrix

We use cross-covariance matrices to study the correlation between selected phenotypes and environmental noise or mutations. Hence, we only consider the phenotype vector’s first 40 out of 200 elements, which comprise the traits subject to selection. We define the Pheno-Cue cross-covariance matrix as

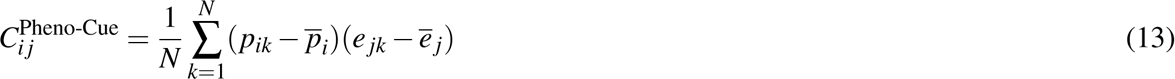

where *p_ik_* is the *i*-th trait of the phenotype vector of the *k*-th individual, *e _jk_* is the *j*-th factor of the environmental cue vector of the *k*-th individual, and *p _j_* and *e _j_*are the population averages of *p_ik_* and *e _jk_*, respectively. We use all 200 elements of the environmental cue vector.

We define the Pheno-Geno cross-covariance matrix as

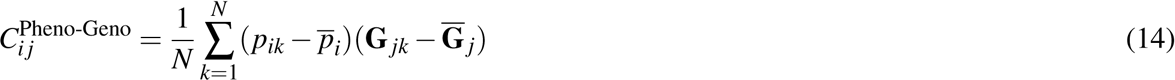

where *p_ik_*and *p_i_* are as defined previously, **G** *_jk_*is the *j*-th element of the vectorized genome of the *k*-th individual, and **G** *_j_* is its population average.

If *C_i j_*is the Pheno-Cue or Pheno-Geno cross-covariance matrix, then the total cross-covariance is defined as

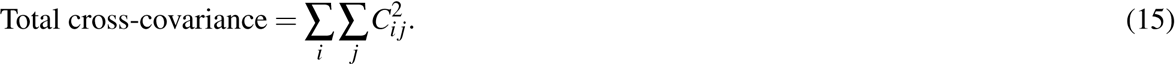

This is used in Figs. 4**b** and 5**b**.

Just as we apply eigenvalue decomposition to a variance-covariance matrix to find principal components that maximize the variance (principal component analysis, PCA), we can apply singular value decomposition (SVD)^50^ to a cross-covariance matrix to find *pairs* of principal components that maximize the cross-covariance^51^. We may apply SVD to any matrix *C* to obtain orthonormal components as follows.

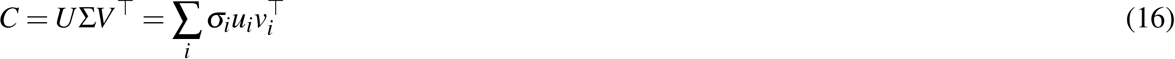

where the superscript *⊤* indicates transpose. In Eq. (16), *u_i_* and *v_i_* are the *i*-th columns of *U* and *V*, respectively, called the *i*-th left and right singular vectors. Σ is a diagonal matrix where the diagonal elements *σ_i_* are singular values arranged in decreasing order. In the case where *C* is a Pheno-Cue (or Pheno- Geno) cross-covariance matrix, the left singular vectors correspond to the principal axes of phenotypic variation in response to the corresponding principal axes (the right singular vectors) of environmental noises (or mutations). The singular values correspond to the cross-covariance between the left and right singular components.

To quantify developmental bias, we used the proportion of the first singular value

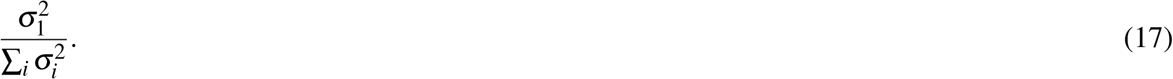

This is used in Figs. 4**c** and 5**c**.

The alignment between the principal axis of phenotypic variation and environmental change is measured by the magnitude of the normalized dot product

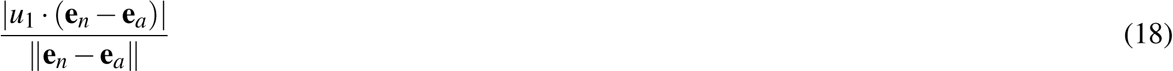

where **e***_n_* is the novel macro-environment and **e***_a_* is the ancestral macro-environment. This is used in Figs. 4**d** and 5**d**.

## Acknowledgments

The authors thank David Marshall and Daphne Teck Ching Lai for their helpful comments and Haziq Jamil for stimulating discussion. E.T.H.N. was supported by the UBD Bursary Award.

## Author contributions

A.R.K. conceived the models and ran preliminary simulations. E.T.H.N. ran large-scale simulations and collected the data. E.T.H.N. and A.R.K. designed the work plan, wrote the computer programs, analyzed the data, and wrote the manuscript.

## Availability of computer code

Computer code is provided in the GitHub repository: https://github.com/arkinjo/evodevo/ tree/Ng23

