## Supplementary Figures for "Plasticity-led evolution as an intrinsic property of developmental gene regulatory networks"

Eden Tian Hwa Ng and Akira R. Kinjo
Department of Mathematics, Faculty of Science,
Universiti Brunei Darussalam, Jalan Tungku Link,
Gadong BE1410, Brunei Darussalam

September 7, 2023

These figures are genotype-phenotype plots of the different models in different novel environments (c.f. Fig. 4 in the main text). See Methods for the definitions of projected phenotypic and genotypic values.

A video of animated trajectories of the simulations is available at
`https://youtu.be/n_cVNgzGG0Y`

### Genotype-Phenotype Plots of the Full model

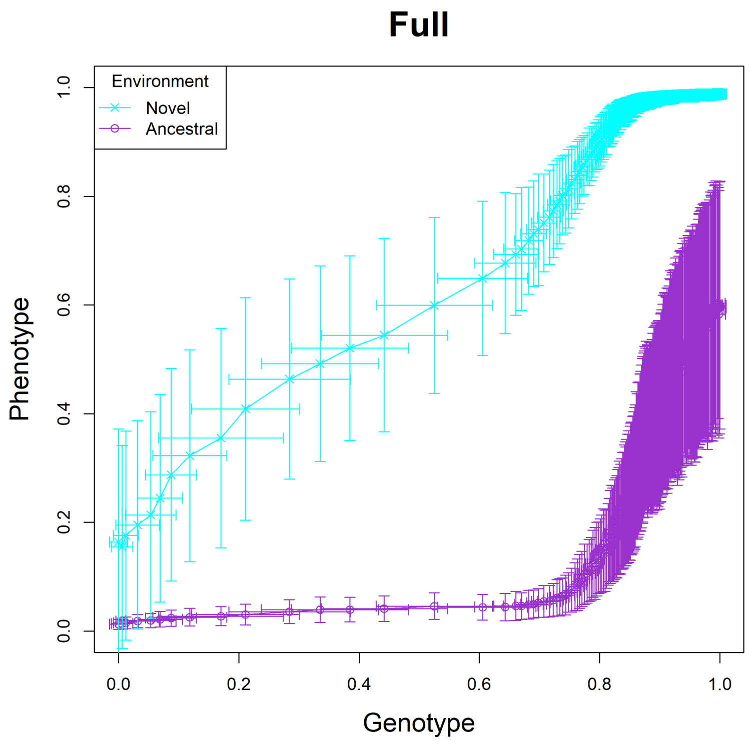


Fig. S1: The Full model, trajectory 01.

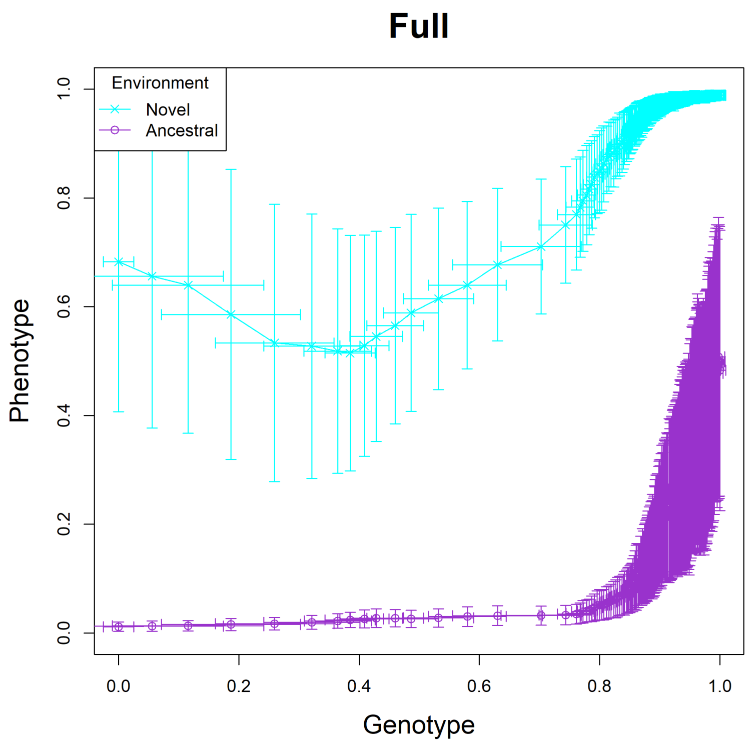


Fig. S2: The Full model, trajectory 02.

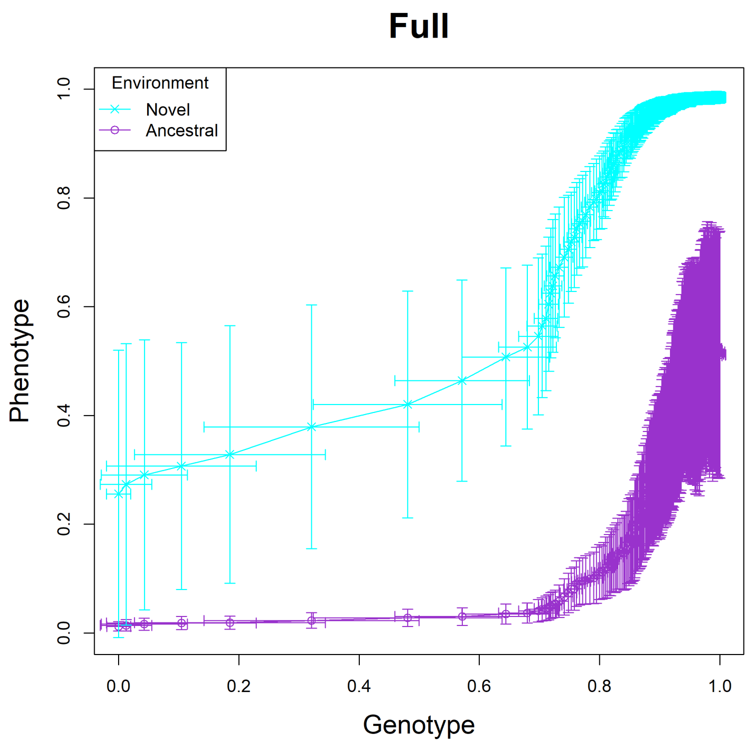


Fig. S3: The Full model, trajectory 03.

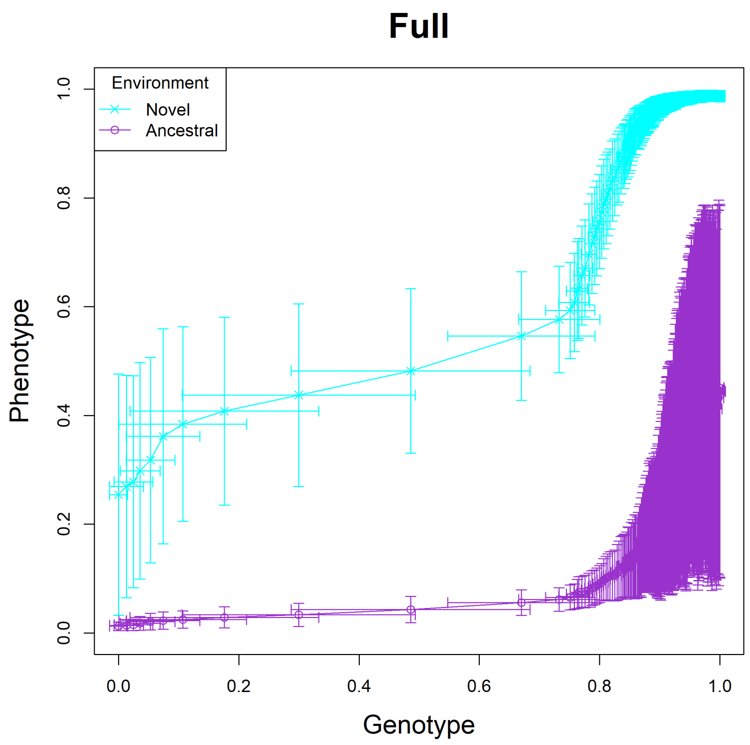


Fig. S4: The Full model, trajectory 04.

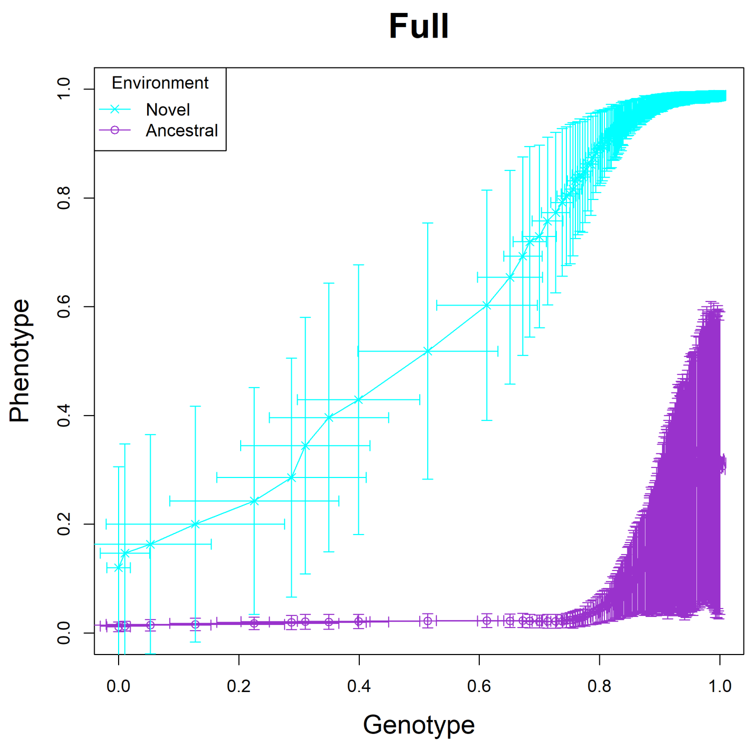


Fig. S5: The Full model, trajectory 05.

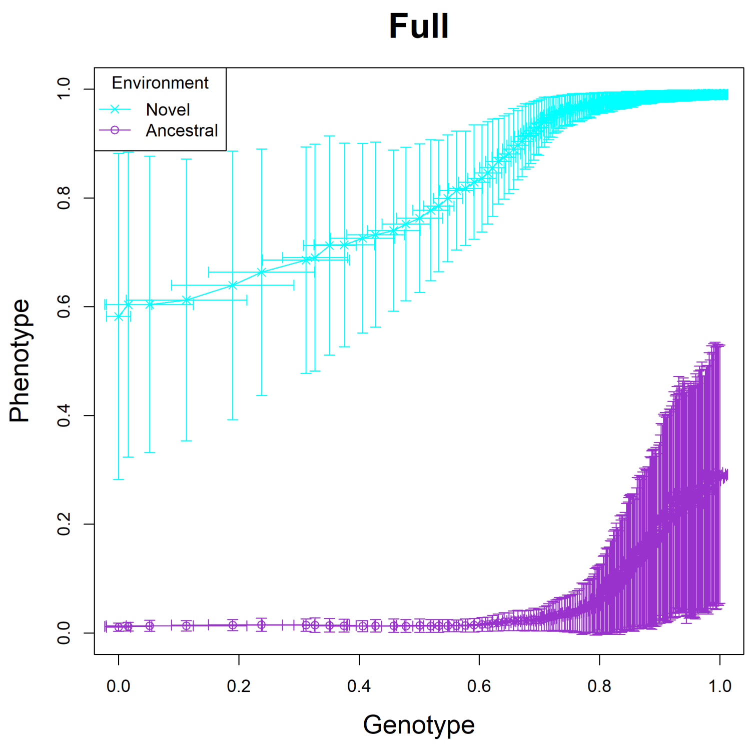


Fig. S6: The Full model, trajectory 06.

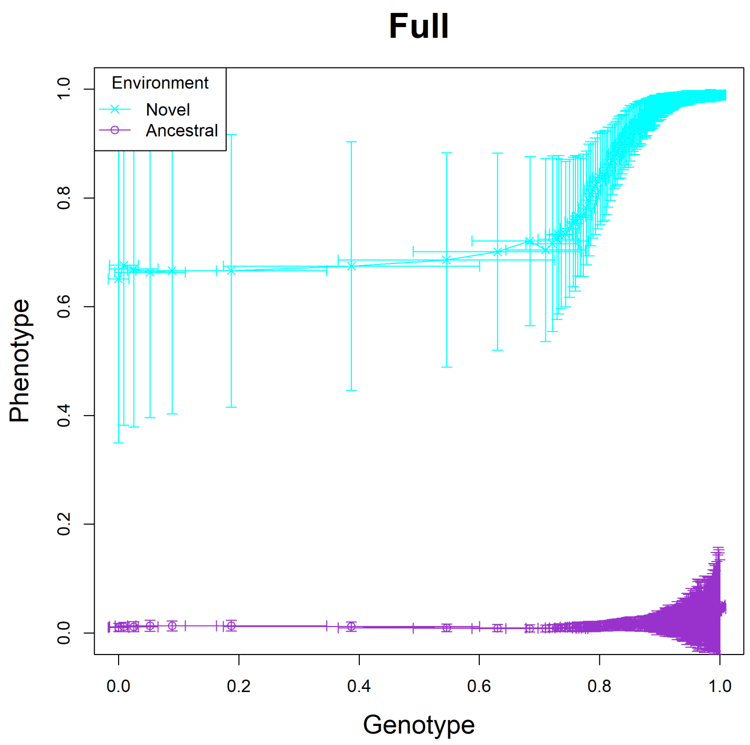


Fig. S7: The Full model, trajectory 07.

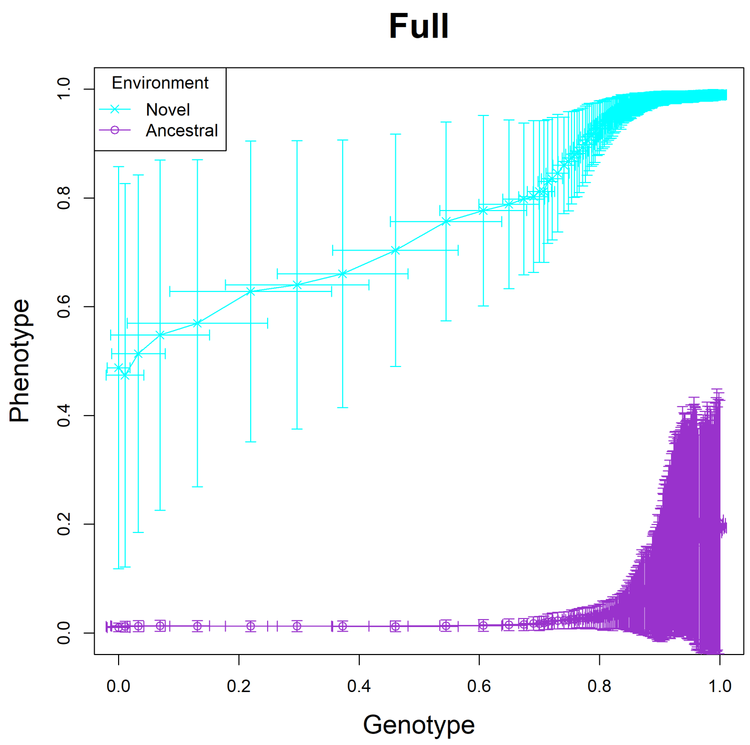


Fig. S8: The Full model, trajectory 08.

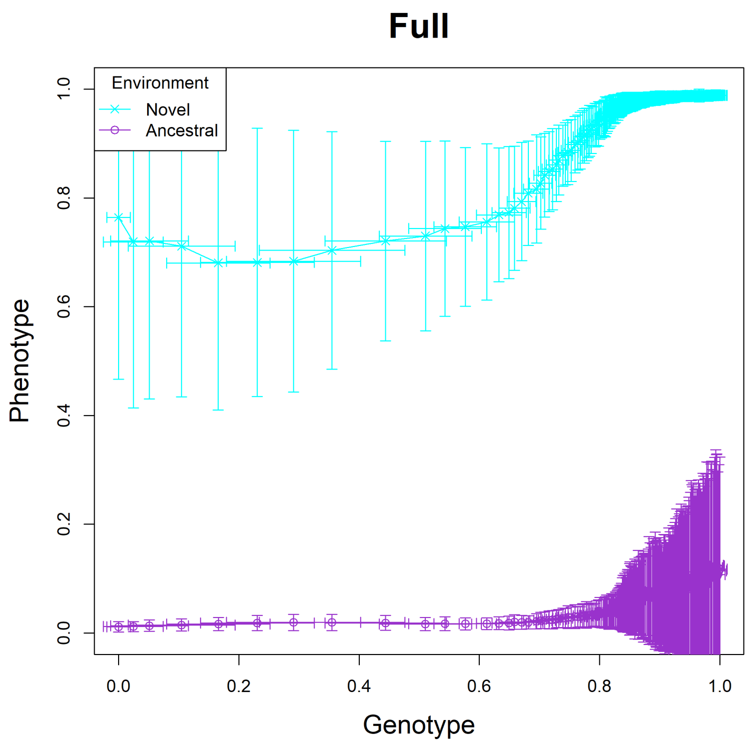


Fig. S9: The Full model, trajectory 09.

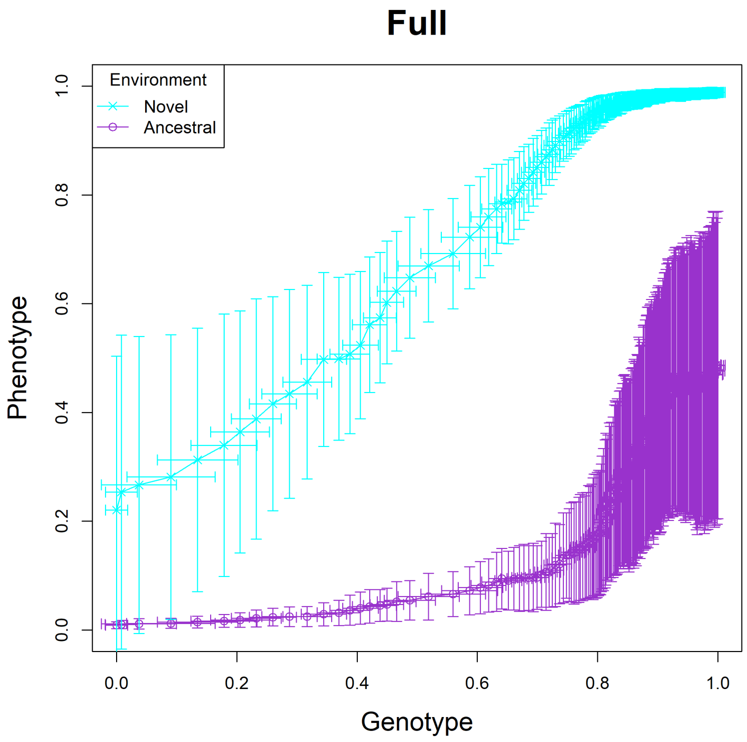


Fig. S10: The Full model, trajectory 10.

### Genotype-Phenotype Plots of the NoHier model

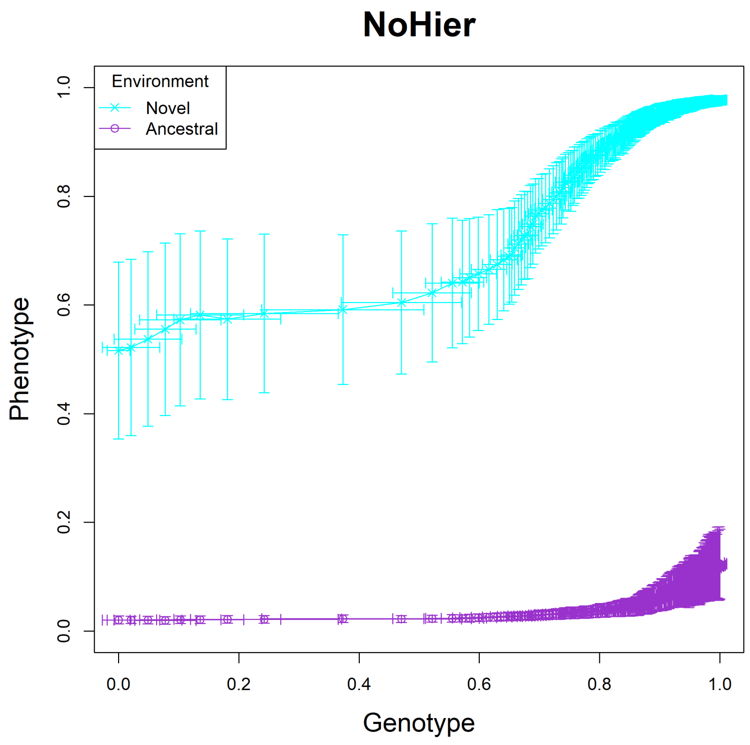


Fig. S11: The NoHier model, trajectory 01.

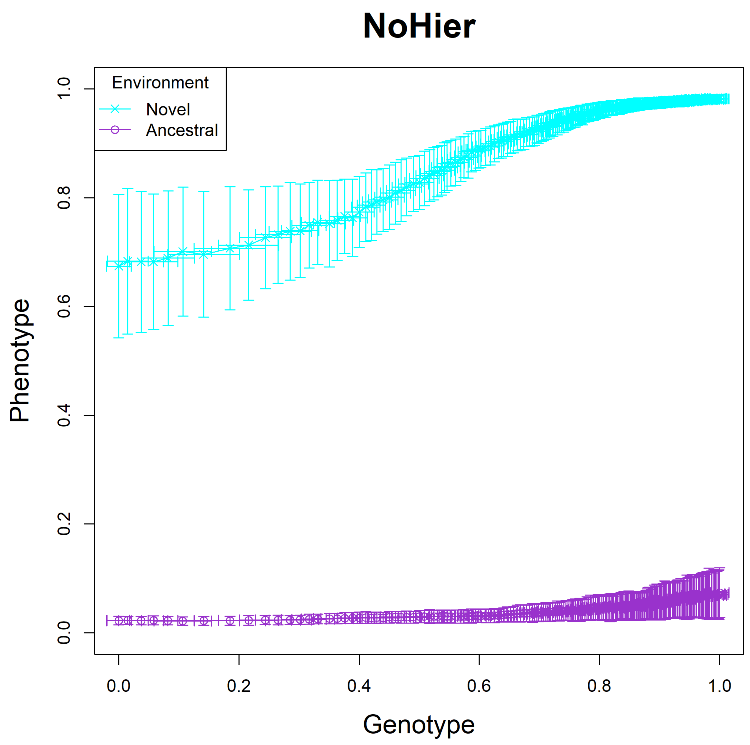


Fig. S12: The NoHier model, trajectory 02.

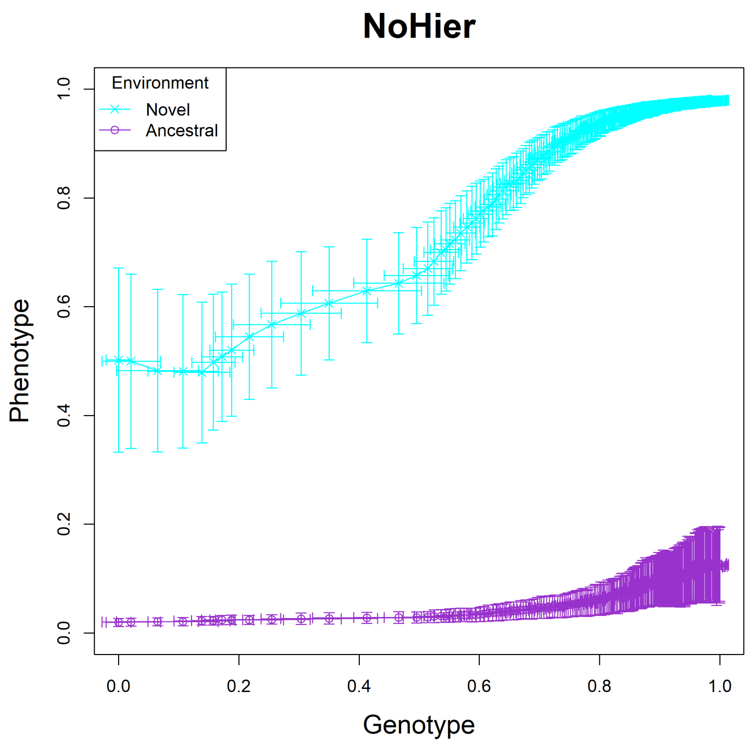


Fig. S13: The NoHier model, trajectory 03.

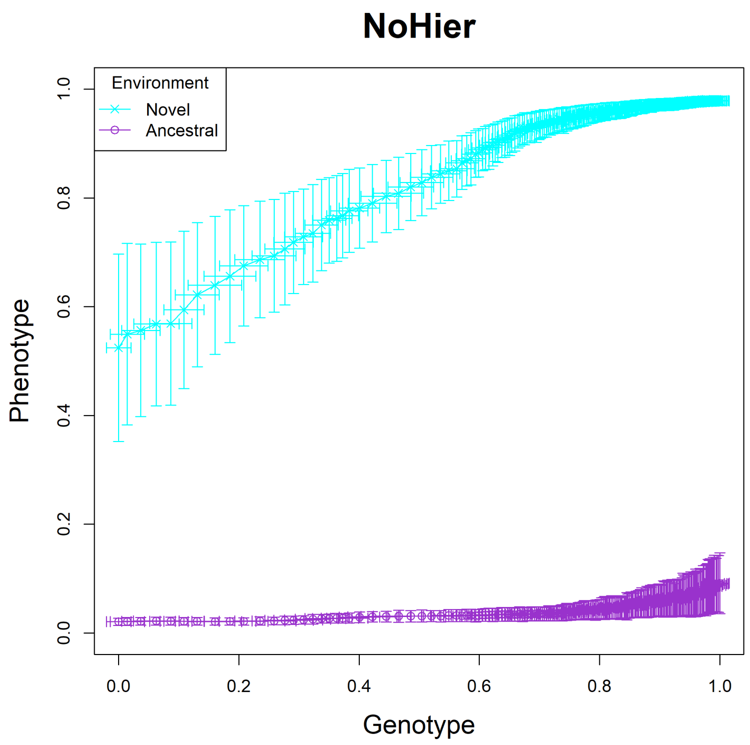


Fig. S14: The NoHier model, trajectory 04.

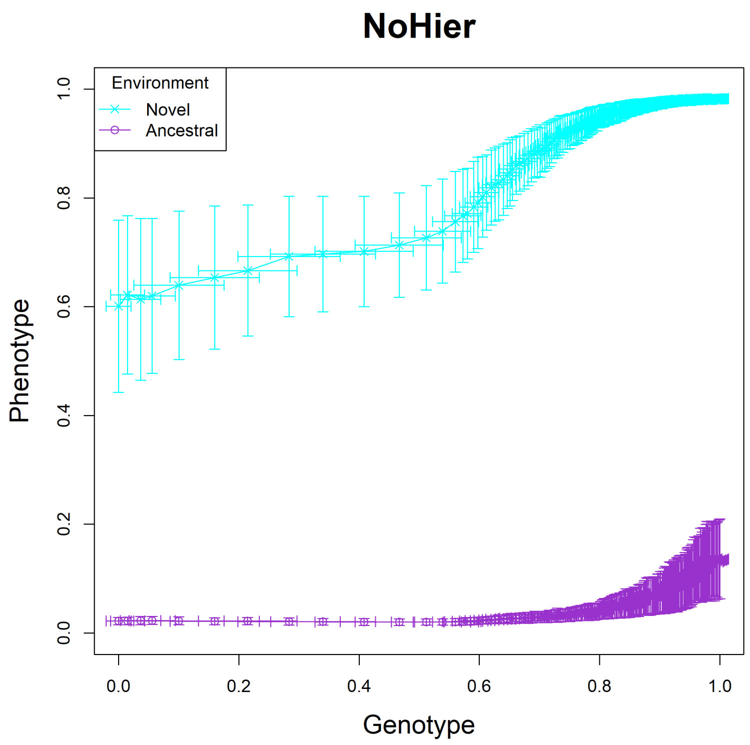


Fig. S15: The NoHier model, trajectory 05.

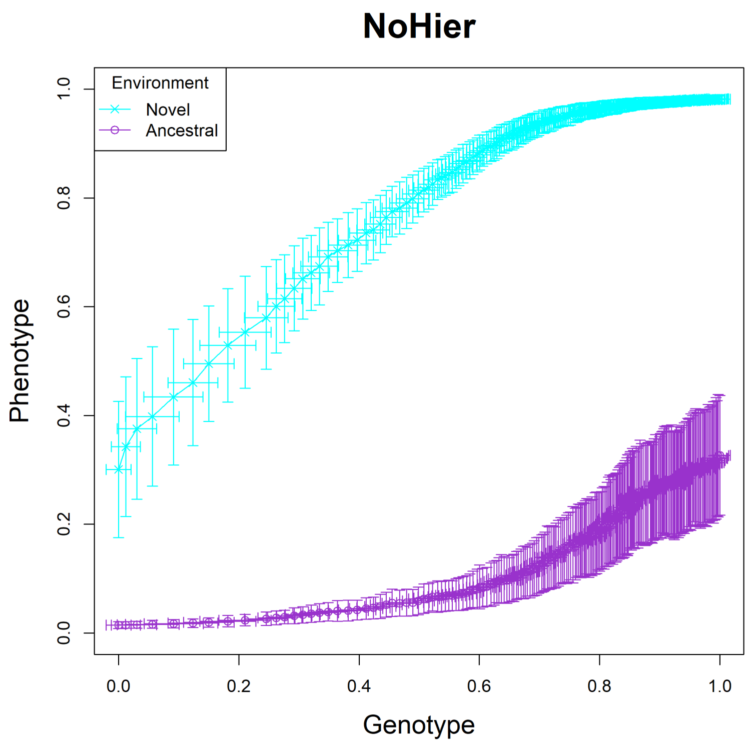


Fig. S16: The NoHier model, trajectory 06.

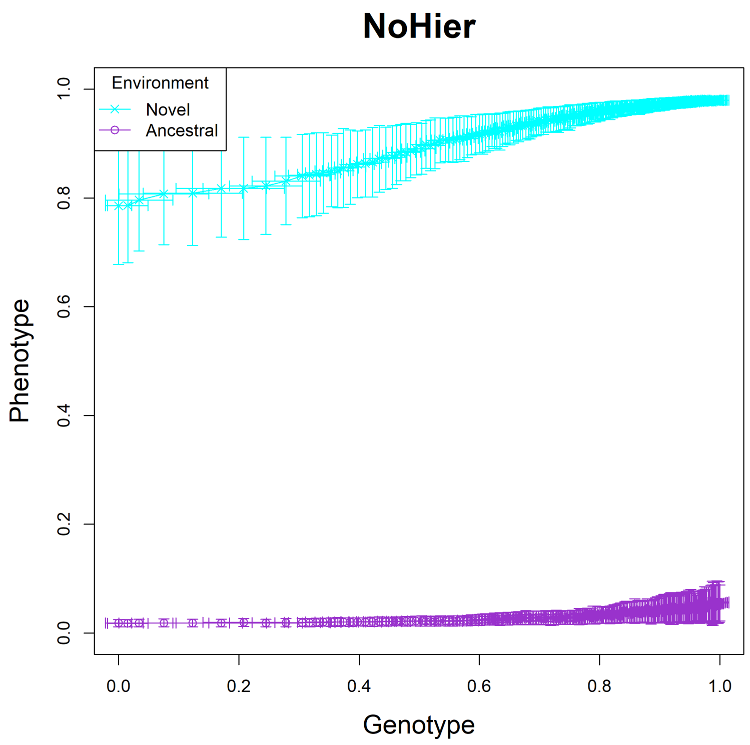


Fig. S17: The NoHier model, trajectory 07.

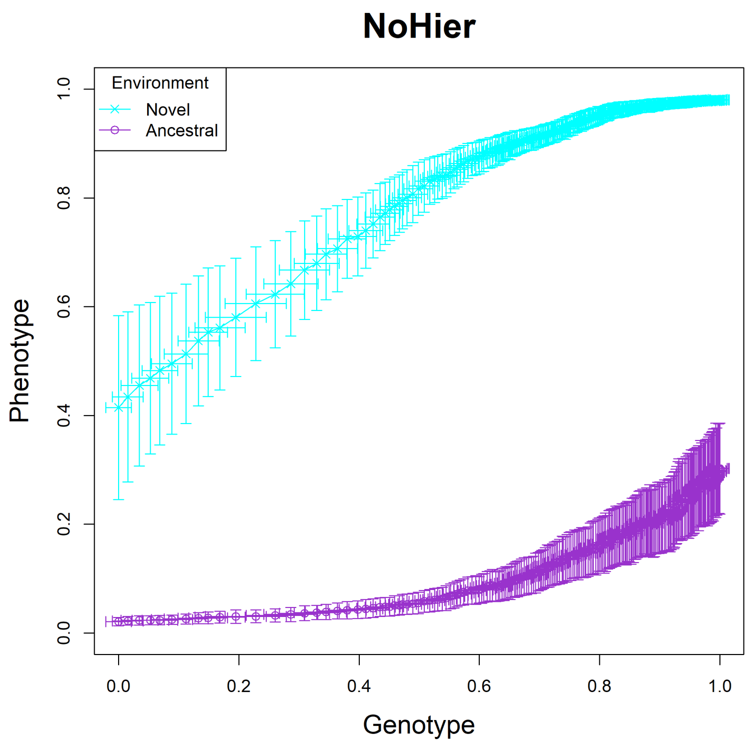


Fig. S18: The NoHier model, trajectory 08.

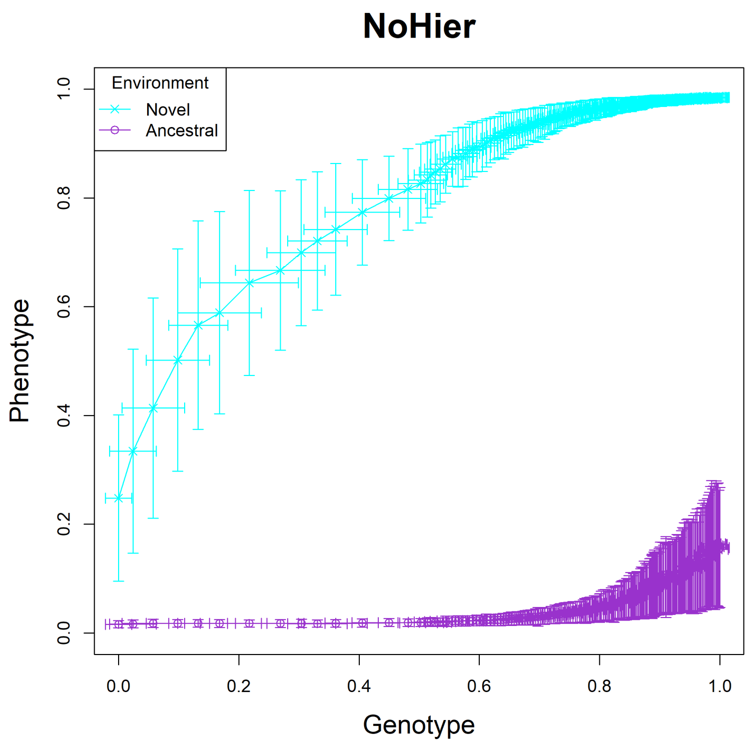


Fig. S19: The NoHier model, trajectory 09.

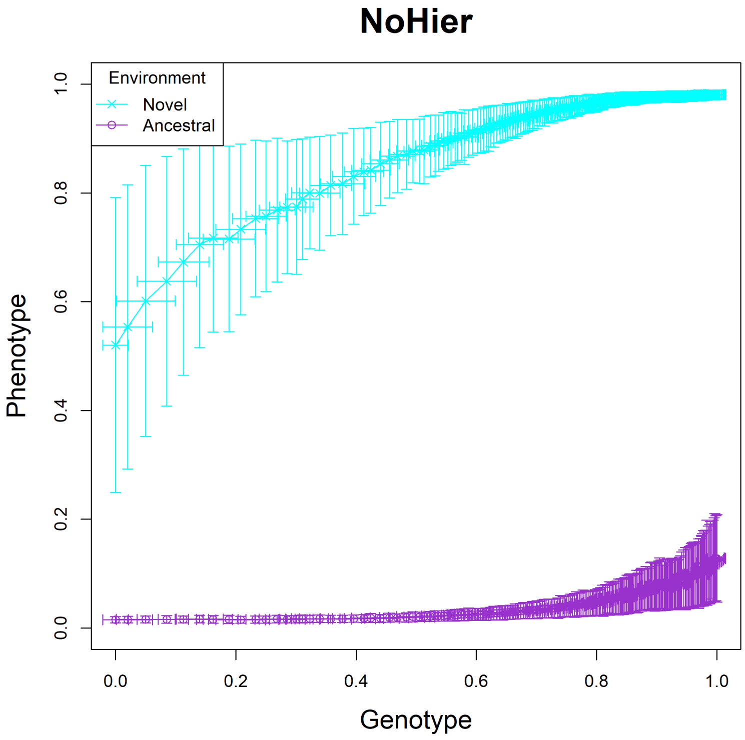


Fig. S20: The NoHier model, trajectory 10.

### Genotype-Phenotype Plots of the NoCue model

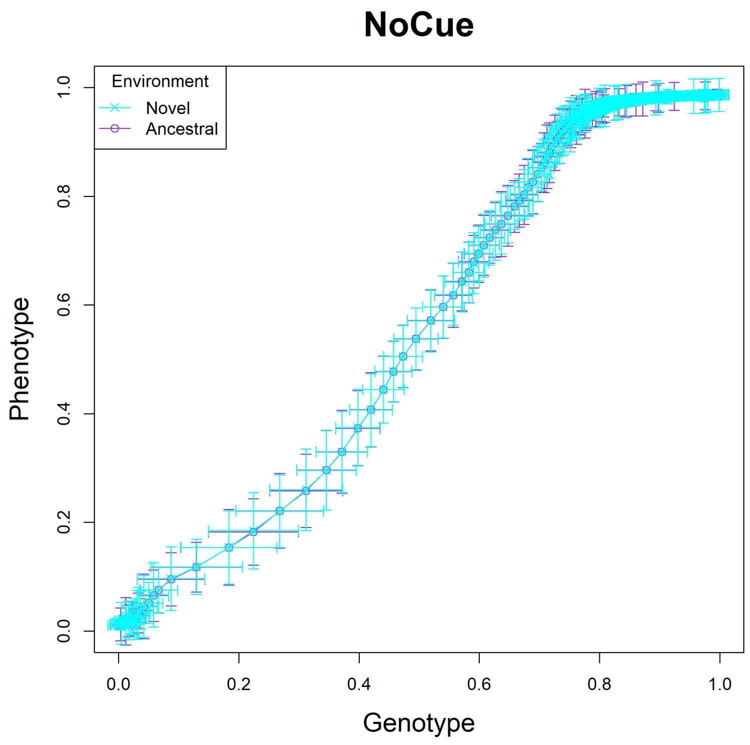


Fig. S21: The NoCue model, trajectory 01.

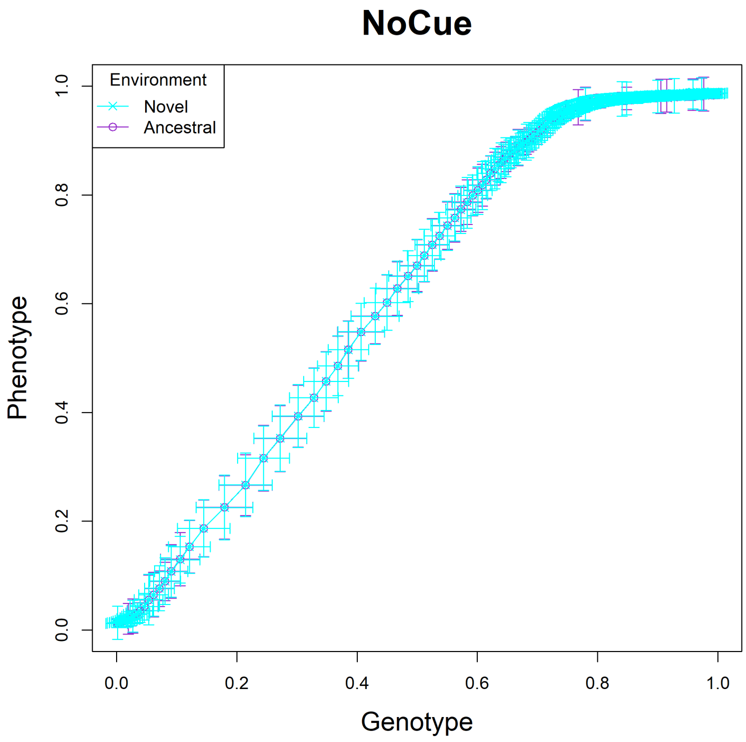


Fig. S22: The NoCue model, trajectory 02.

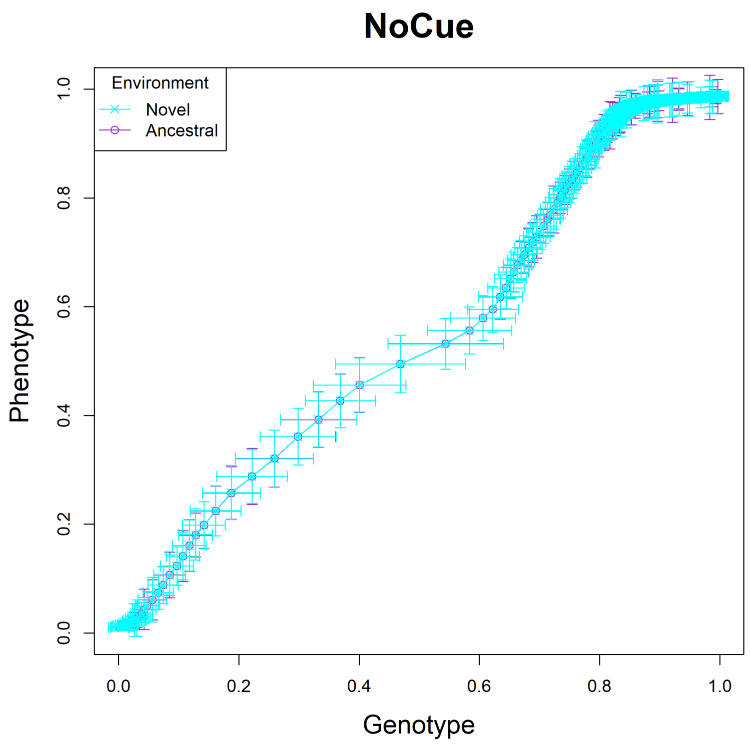


Fig. S23: The NoCue model, trajectory 03.

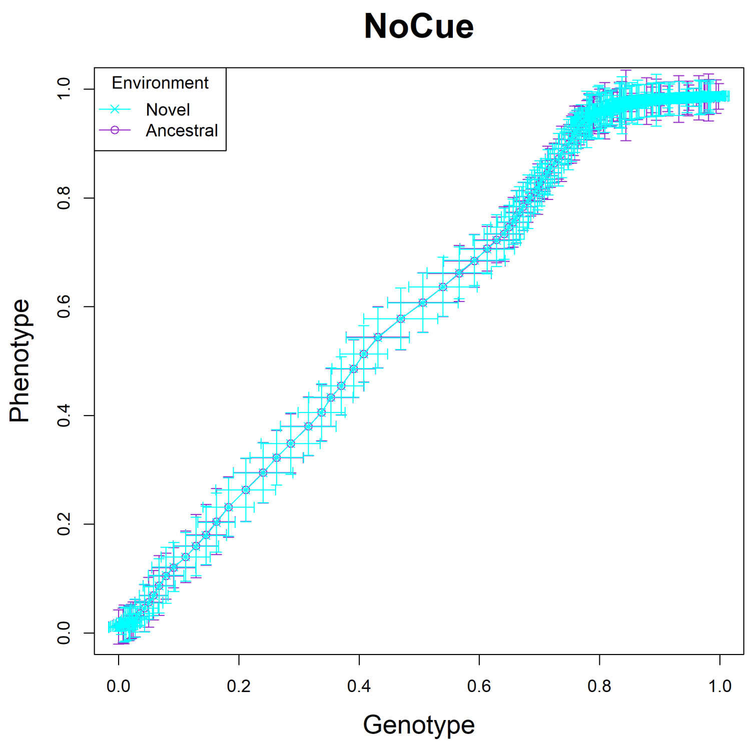


Fig. S24: The NoCue model, trajectory 04.

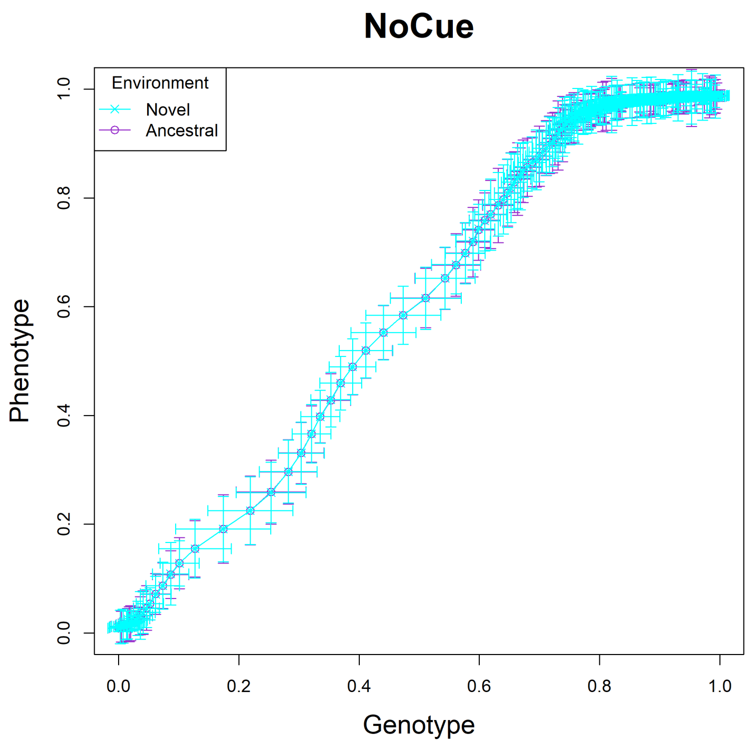


Fig. S25: The NoCue model, trajectory 05.

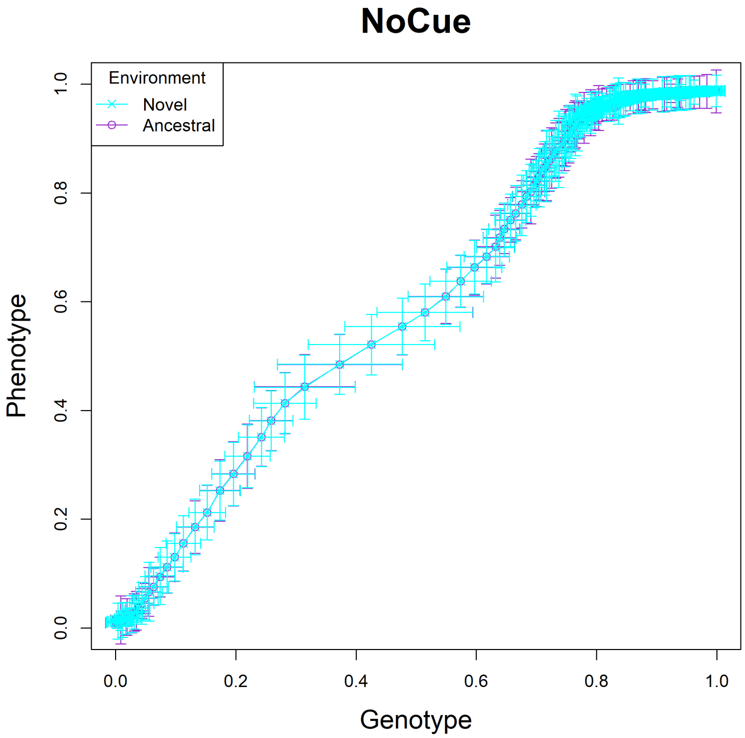


Fig. S26: The NoCue model, trajectory 06.

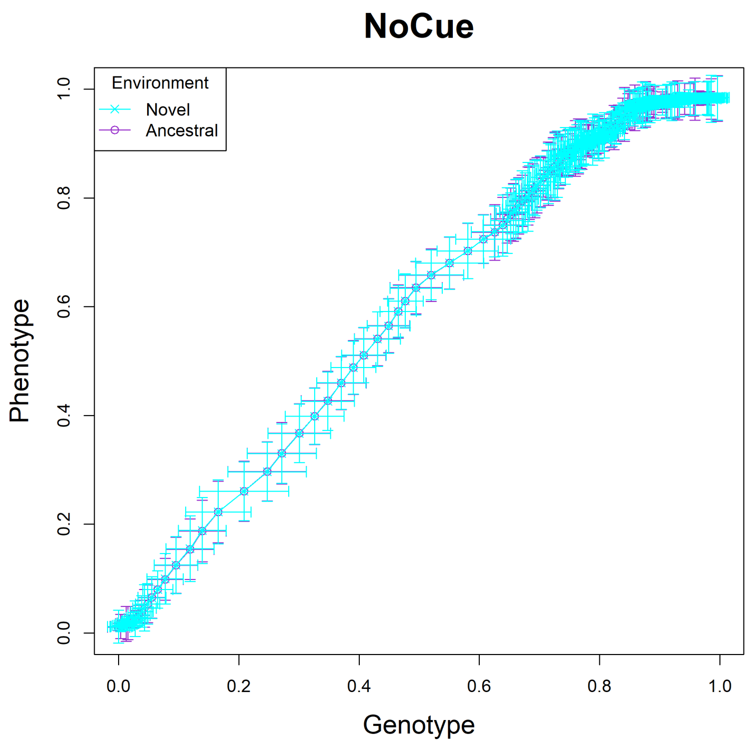


Fig. S27: The NoCue model, trajectory 07.

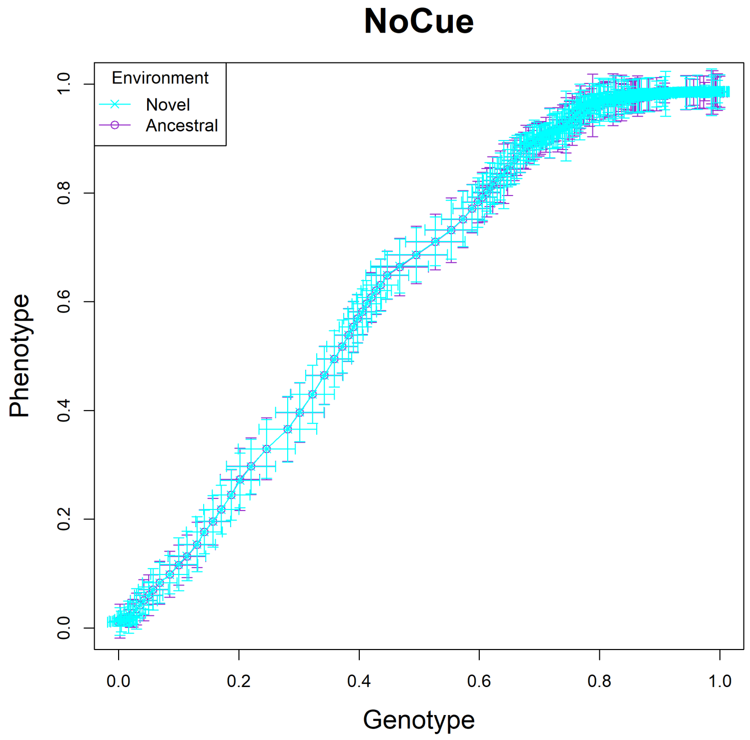


Fig. S28: The NoCue model, trajectory 08.

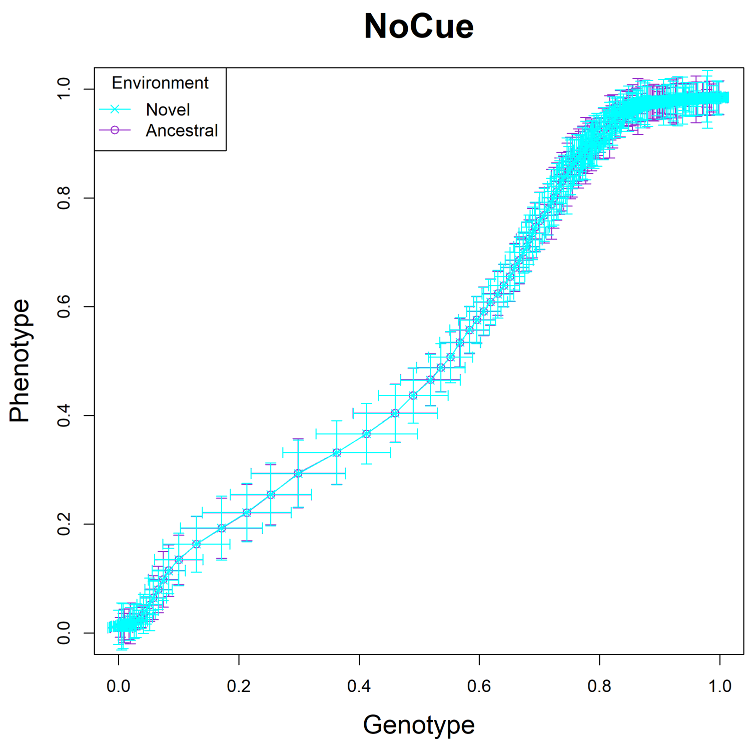


Fig. S29: The NoCue model, trajectory 09.

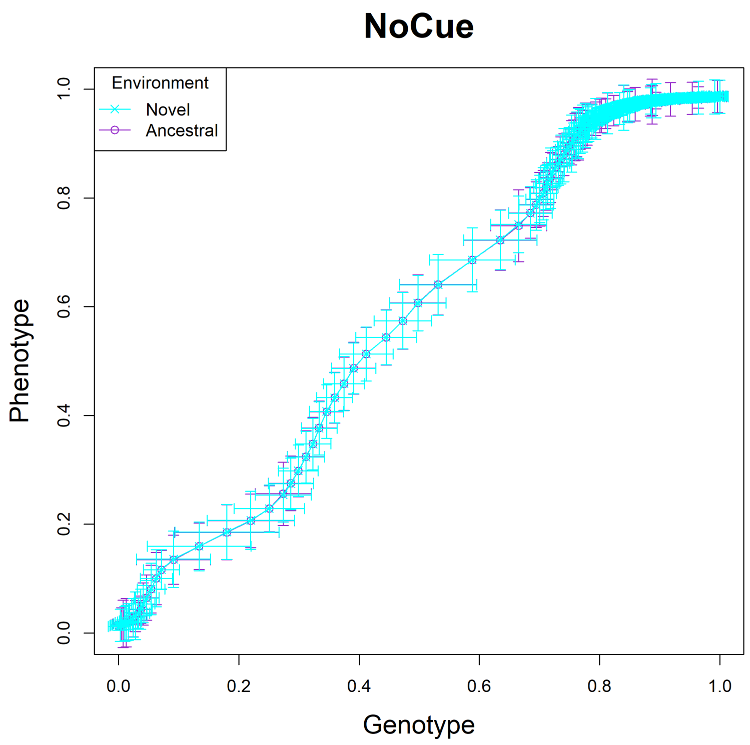


Fig. S30: The NoCue model, trajectory 10.

### Genotype-Phenotype Plots of the NoDev model

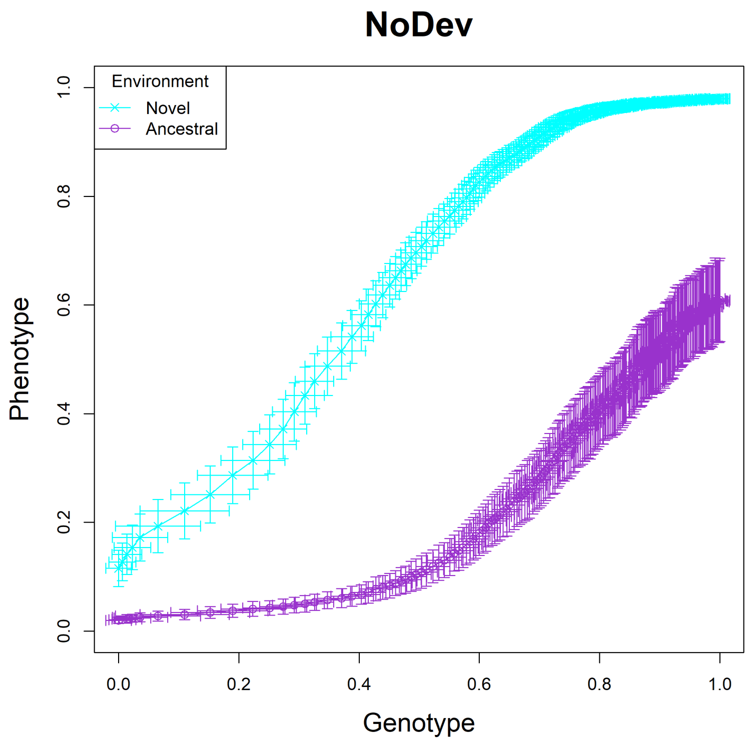


Fig. S31: The NoDev model, trajectory 01.

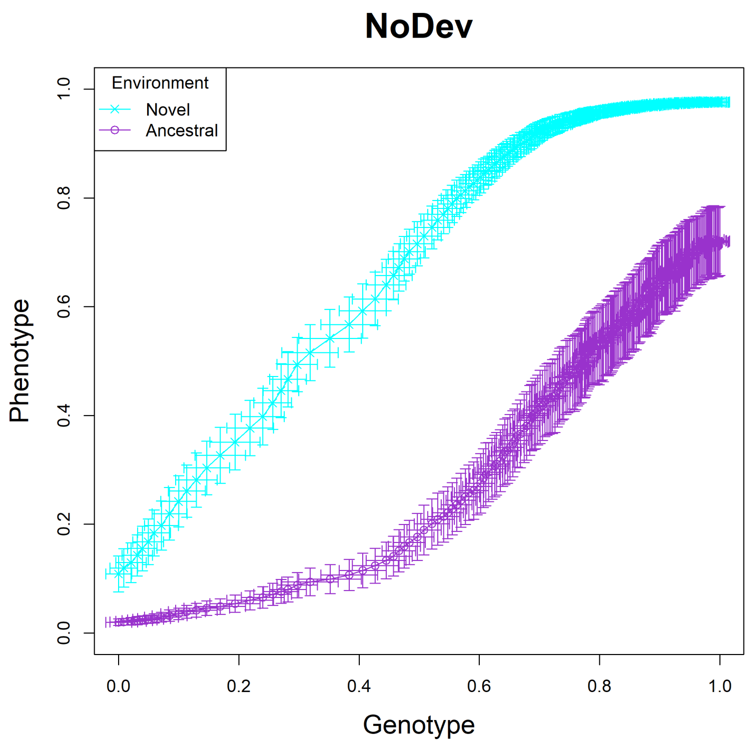


Fig. S32: The NoDev model, trajectory 02.

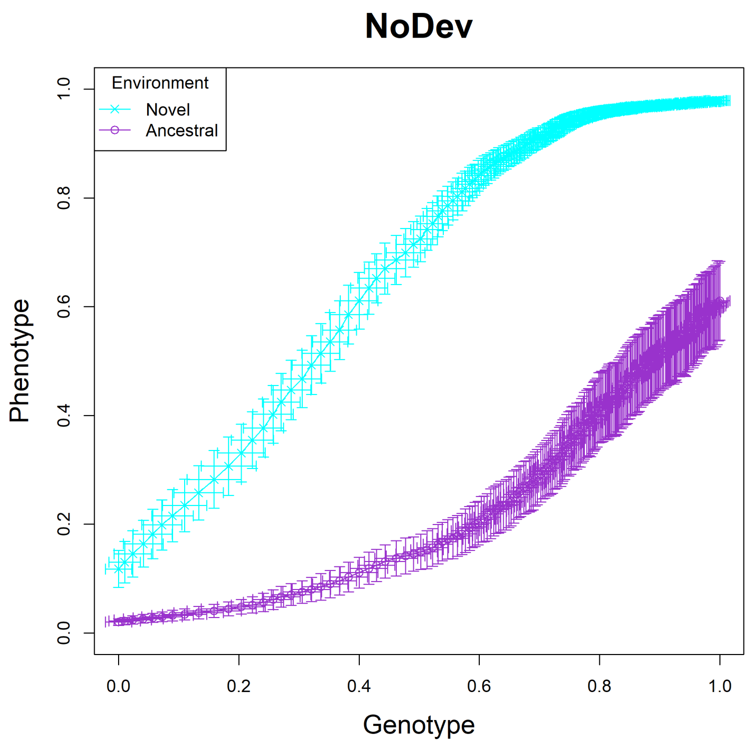


Fig. S33: The NoDev model, trajectory 03.

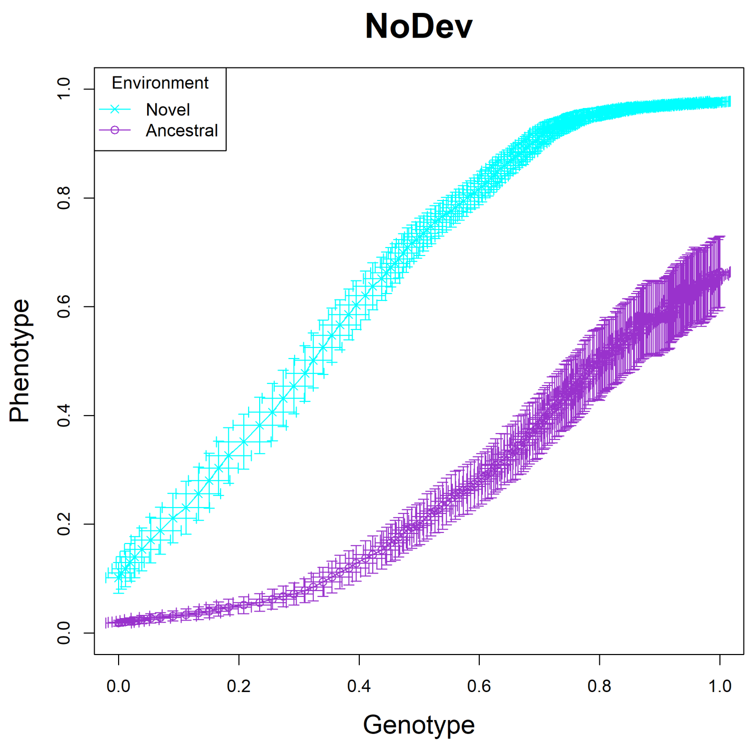


Fig. S34: The NoDev model, trajectory 04.

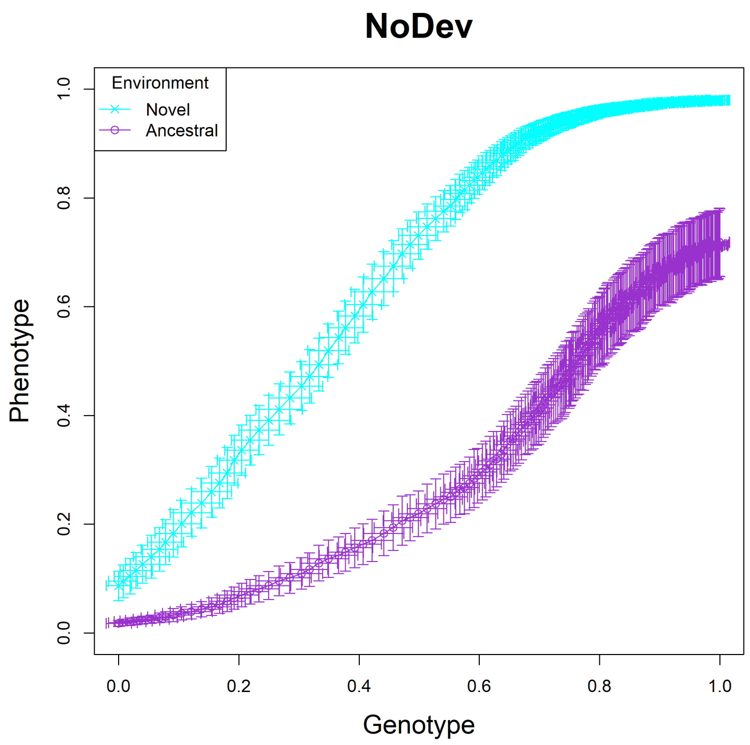


Fig. S35: The NoDev model, trajectory 05.

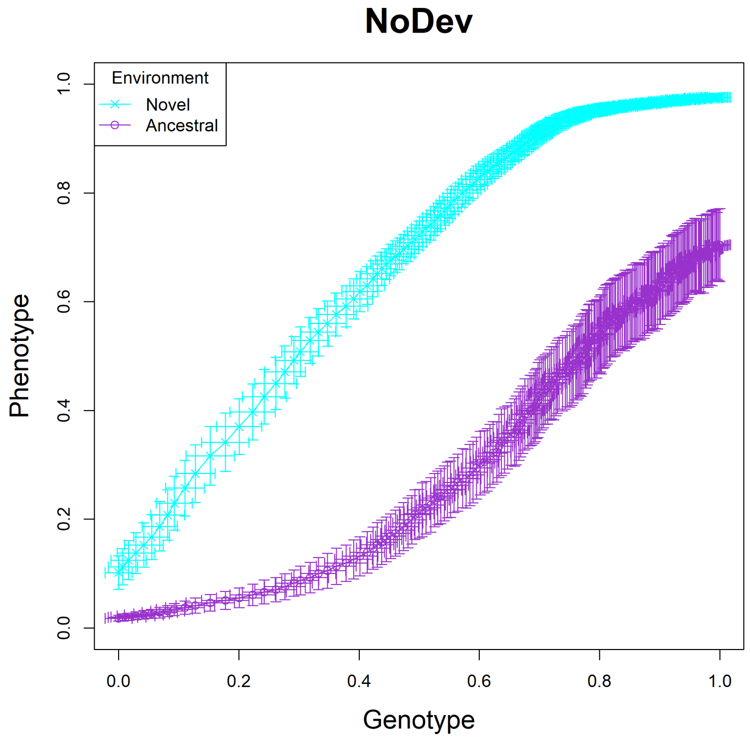


Fig. S36: The NoDev model, trajectory 06.

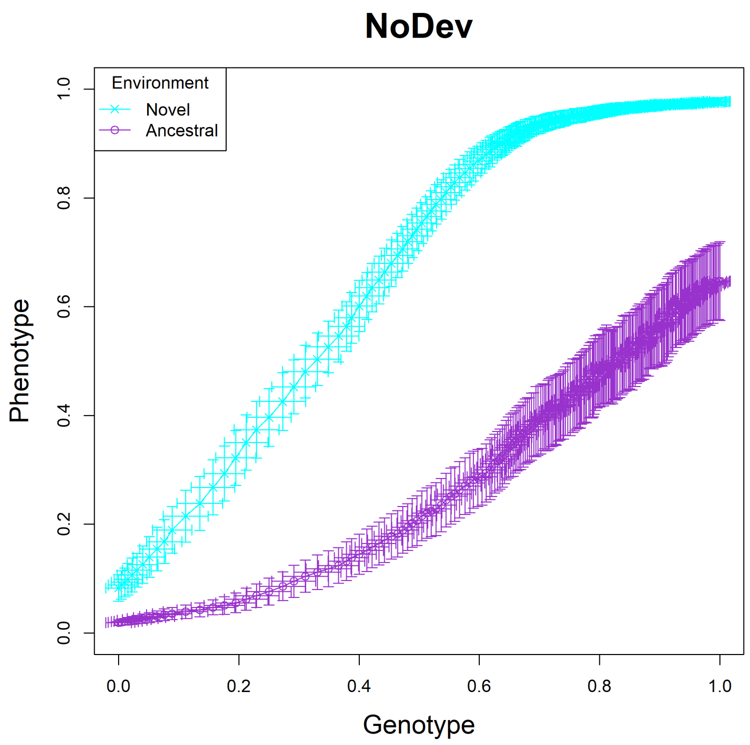


Fig. S37: The NoDev model, trajectory 07.

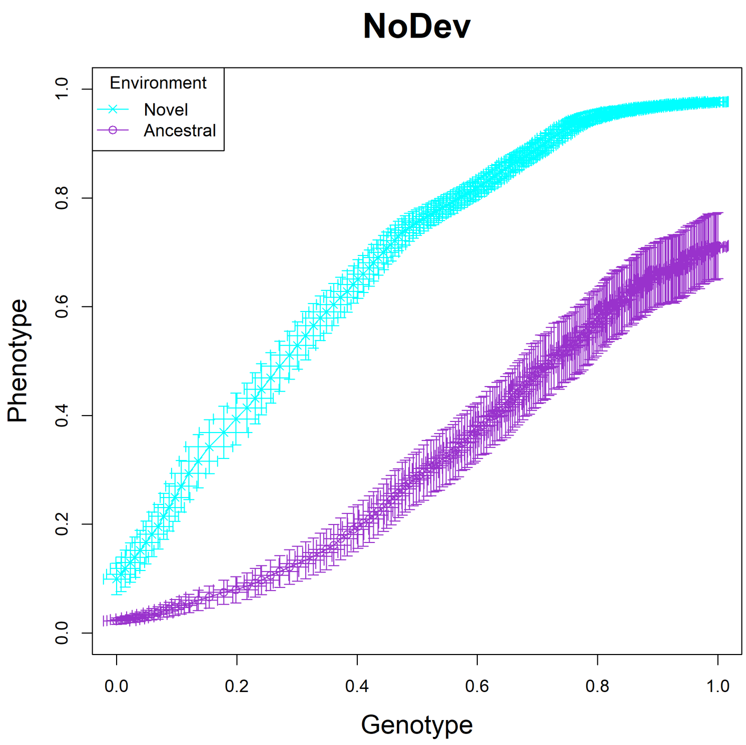


Fig. S38: The NoDev model, trajectory 08.

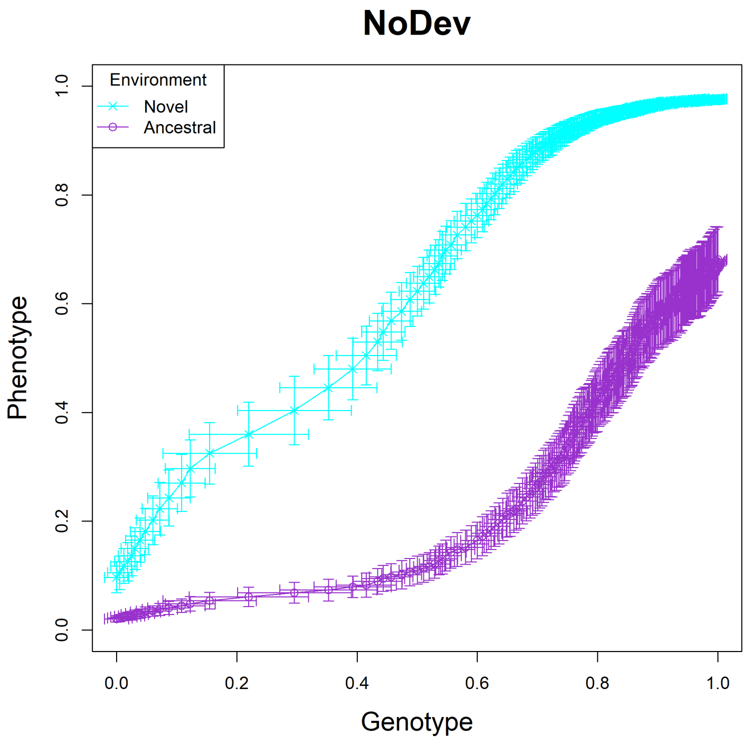


Fig. S39: The NoDev model, trajectory 09.

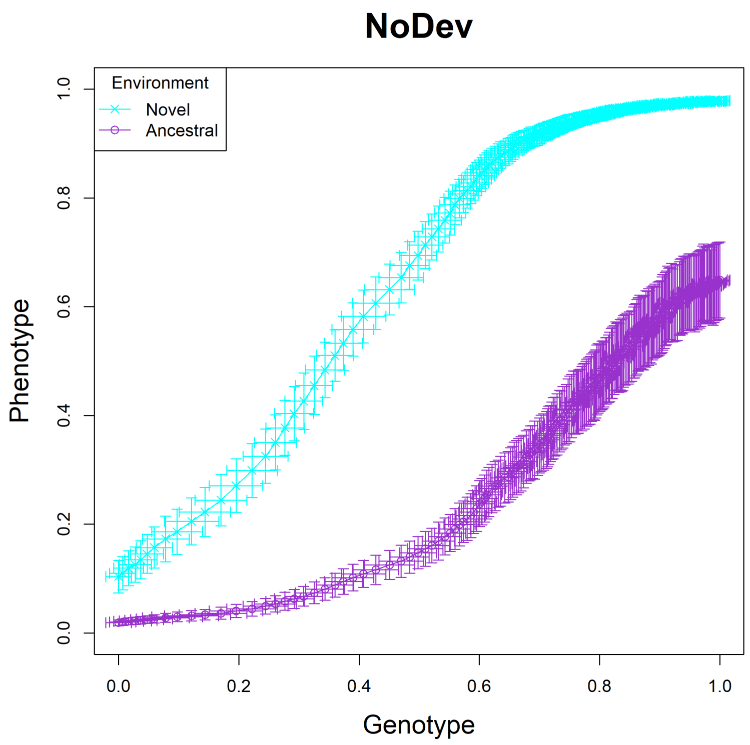


Fig. S40: The NoDev model, trajectory 10.
